# DNA-Programmed Condensate-Membrane Wetting and Cellular Internalization

**DOI:** 10.64898/2026.07.15.738578

**Authors:** Zeyu Chen, Weifen Chen, Jingyi Ye, Dan Lu, Markita P. Landry, Honglu Zhang, Chunhai Fan, Huan Zhang

## Abstract

Deformable condensates offer dynamic interfaces for biomolecule delivery, yet membrane adhesion does not necessarily lead to cellular internalization. The physical transition that determines whether a membrane-bound soft material remains surface-anchored or proceeds through wetting towards productive uptake remains poorly understood, particularly at active living-cell membranes. Here, we engineer sequence-defined DNA condensates through liquid-liquid phase separation (LLPS) and program their interfacial behavior by tuning sticky-end valency and cholesterol organization. These molecular designs precisely regulate condensate fluidity, fusion dynamics and internal organization, generating distinct states of weak contact, persistent anchoring and rapid wetting. Increasing cholesterol-mediated affinity does not enhance uptake. Instead, productive internalization emerges from a balance between membrane adhesion and condensate fluidity and deformability. Native membrane composition further modulates condensate interfacial fate across mammalian cells and plant protoplasts. DNA condensates enrich and deliver CpG ODNs, mRNA (∼2000 nt) and proteins, while cargo loading experiments reveal that preserving condensate architecture is essential for functional delivery. Our findings identify wetting competence as a design parameter for controlling soft material engagement and cellular entry.

## Introduction

Biomolecular condensates are dynamic, deformable materials formed through liquid-liquid phase separation (LLPS) of proteins, nucleic acids and other multivalent components^1–3^. In living cells, they compartmentalize biomolecules and regulate processes ranging from transcription and signaling to autophagy and immune responses^4–8^. Their all-aqueous interiors, molecular exchange dynamics and capacity to recruit diverse macromolecules have also inspired synthetic condensates as cell-mimetic compartments, reaction microenvironments and biomolecular delivery systems^9, 10^. Unlike conventional solid particles, however, condensates can reorganize and deform upon contact with biological interfaces. This raises the fundamental materials question of whether a deformable condensate remains surface-bound, spreads along the membrane or proceeds towards cellular internalization after reaching and adhering to the cell membrane.

Cellular delivery systems are commonly optimized in terms of cargo loading, colloidal stability, surface charge, membrane affinity and receptor-mediated uptake^11, 12^. Much less attention has been given to the physical transition that connects initial membrane adhesion to efficient internalization. For deformable condensates, this transition may involve interfacial reorganization and membrane wetting, which expand the condensate-membrane contact area and couple condensate deformation to membrane remodelling^13, 14^. Reconstituted biomimetic systems, particularly giant unilamellar vesicles (GUVs), have provided a quantitative framework for describing condensate-membrane wetting through contact-angle geometry, interfacial tension and membrane deformation^14–16^. These studies show that membrane association alone does not necessarily lead to spreading, wrapping or engulfment, because the resulting interfacial fate is jointly governed by adhesion strength, condensate interfacial tension, membrane tension and bending elasticity^14–16^.

Whether these principles can be translated into design rules for living cell interfaces remains unclear. Native plasma membranes are not passive equilibrium substrates, but compositionally heterogeneous and mechanically dynamic boundaries containing diverse lipids, cholesterol-rich domains, membrane proteins, cytoskeletal constraints and active endocytic machineries^17^. Thus, strong membrane affinity alone does not guarantee cellular entry, productive internalization requires a transition from adhesion to wetting, enabled by appropriate molecular mobility and interfacial deformability. Understanding how membrane adhesion progresses to wetting and cellular internalization would provide a physical basis for designing soft materials with distinct cellular fates, ranging from persistent surface anchoring for localized membrane activity to efficient internalization for intracellular delivery.

DNA nanotechnology provides a precisely controllable platform for addressing this problem. Predictable base-pairing and chemical addressability allow molecular connectivity, interaction valency and functional-group positioning to be programmed with high precision^18^. Sticky-end-mediated assembly can drive DNA nanostructures to undergo LLPS and form micron-scale droplets whose phase behavior, molecular mobility and internal organization can be tuned through sequence design^19–24^. Prior studies have established the programmability of DNA condensate formation and dynamics, including sequence-dependent fusion, fission and molecular recruitment^20, 22, 25^. However, it remains largely unexplored how molecular architecture controls the interfacial fate of preformed DNA condensates at living-cell membranes, and in particular whether membrane adhesion progresses towards persistent anchoring, productive wetting or cellular internalization.

Here, we engineer DNA-programmed condensates to study how molecular architecture and material state control interfacial fate at active living cell membranes. Multibranched DNA monomers were designed to undergo LLPS through sticky-end hybridization, while cholesterol anchors were introduced with defined valency and spatial arrangement. By varying sticky-end valency together with cholesterol valency and positioning, we programmed condensate fluidity, fusion dynamics and internal organization. These material states generated three distinct membrane-interaction behaviors, weak contact, stable surface anchoring and rapid wetting, and revealed that membrane affinity alone is insufficient to promote cellular uptake. Instead, efficient internalization was associated with an appropriate balance between membrane adhesion and condensate deformability. Experiments across mammalian cells and compositionally distinct plant protoplasts further showed that native membrane composition selects condensate interfacial fate. Finally, delivery of CpG oligodeoxynucleotides (ODNs), mRNA and proteins demonstrated the functional consequences of this design rule, indicating that cargo performance depends not only on loading but also on preservation of a wetting-competent condensate state. This work thus establishes a molecular design principle for directing the interfacial fate of deformable condensates, from persistent surface anchoring to productive wetting and cellular internalization, with direct implications for biomolecule delivery.

**Scheme 1.**
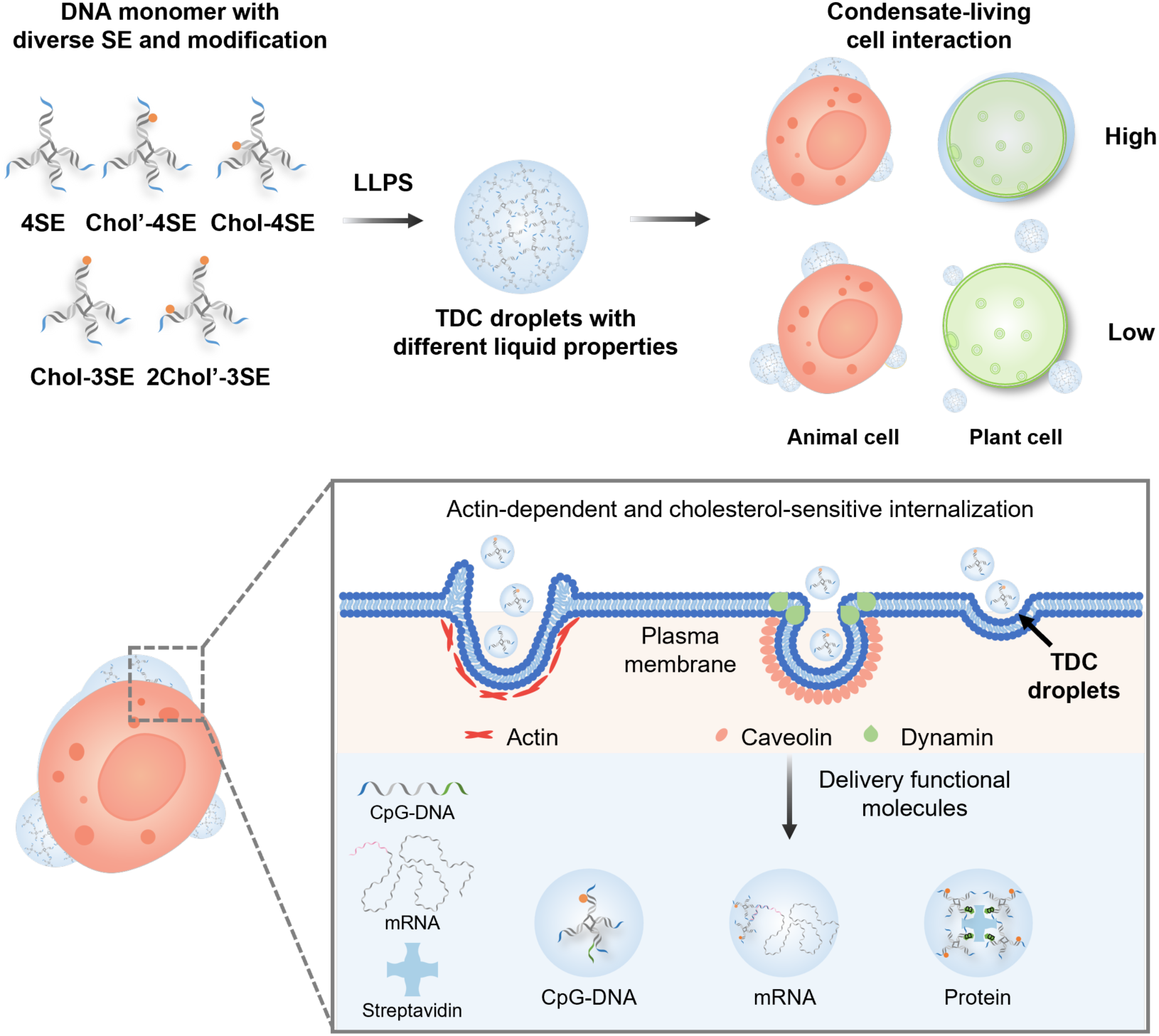
DNA-programmed condensate droplets direct membrane wetting, anchoring and internalization for biomolecule delivery. Sequence-defined DNA monomers with tunable sticky-end valency and cholesterol modification undergo LLPS to form condensate droplets with programmable material properties. These droplets engage living cell membranes through defined interfacial states, including weak contact, membrane wetting and surface anchoring, which shape subsequent internalization pathways involving actin remodeling and cholesterol-sensitive membrane domains. The same DNA condensate platform enables enrichment and intracellular delivery of CpG ODNs, mRNA and protein cargoes.

## Results and Discussion

### Molecular architecture programs DNA condensate material states

To establish a programmable material platform for investigating condensate-membrane interactions, we first constructed a library of tetrapod DNA condensates (TDCs) in which sticky-end valency and the number and spatial organization of cholesterol moieties were systematically varied. A series of multibranched tetrapod DNA frameworks were designed as basic monomers to form condensate droplets through phase separation driven by sticky-end (SE) interactions. Each monomer was assembled from four or five single-stranded DNA sequences (Table S1), with each branch containing a 16-base-pair (16-bp) duplex and terminating in a 4-nucleotide (4-nt) palindromic sticky end^23^. To tune hydrophobicity and molecular organization, cholesterol groups were introduced either adjacent to or distal from the SEs, and at different valencies. When incubated in PBS at 37 °C, sticky-end hybridization drove tetrapod DNA monomers to undergo LLPS, forming micron-scale tetrapod DNA condensates (TDCs) within one hour (Fig. 1a).

**Fig. 1.**
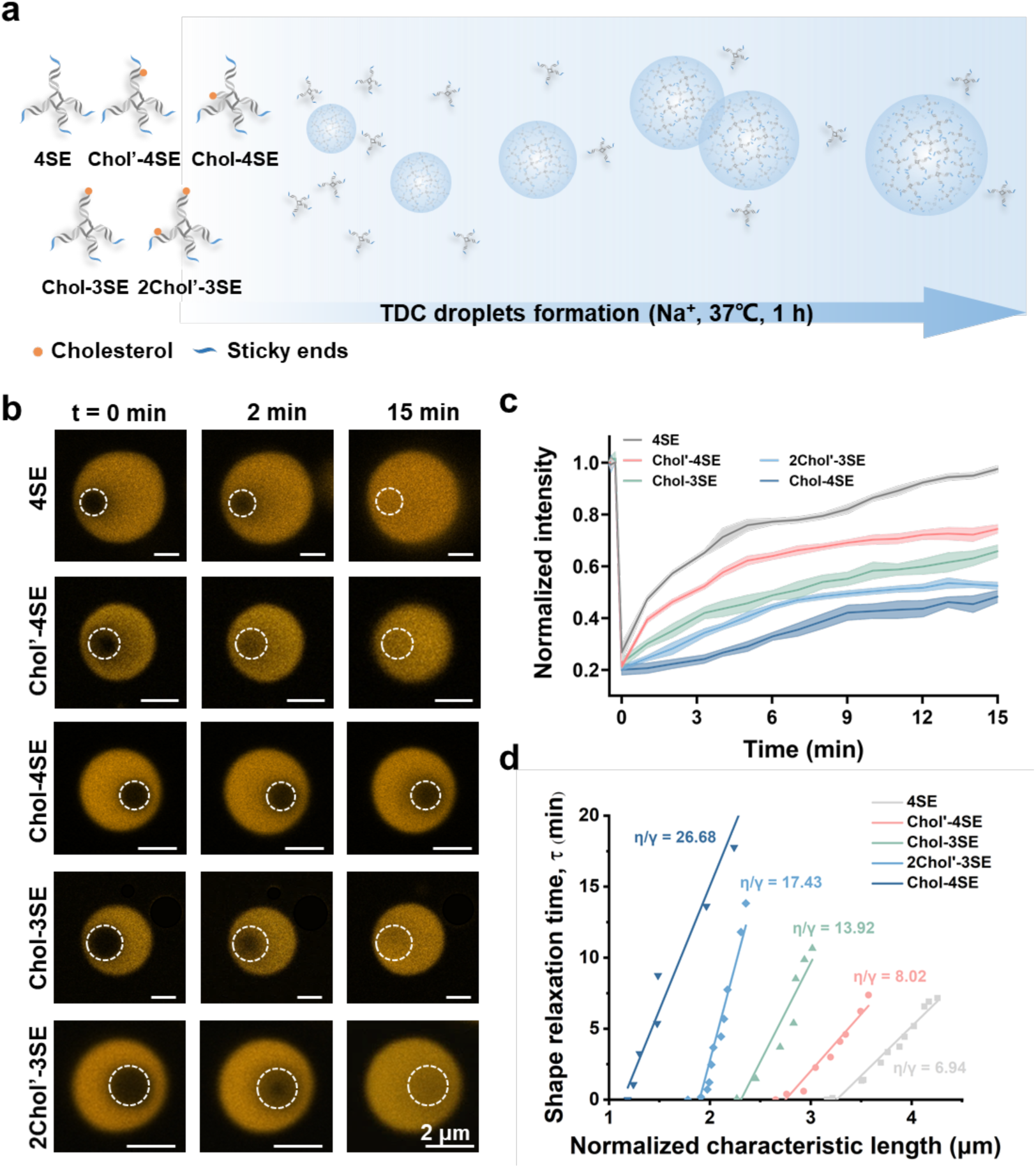
Formation and dynamics of DNA-programmed condensate droplets. (a) Schematic illustration of TDC formation from multibranched DNA monomers with different sticky-end valencies and cholesterol modification patterns. (b) Representative CLSM images of partially photobleached Cy3-labeled TDC droplets at indicated time points. Dashed circles indicate the photobleached regions. Scale bars: 2 μm. (c) Normalized fluorescence recovery curves showing differential molecular mobility among TDC droplets. Solid lines and shaded regions represent the mean and s.d., respectively (n=3). (d) Fusion dynamics of TDC droplets quantified by shape relaxation analysis. The slope of the linear fit corresponds to η/γ, the ratio of apparent viscosity to interfacial tension. Relaxation time (τ) is plotted against characteristic length (*L)* and the slope of each linear fit yields η/γ, the ratio of apparent viscosity to interfacial tension (n=1).

For clarity, the TDCs were named according to the number of sticky ends and cholesterol (Chol) modifications (Fig. 1a). The unmodified four-branched monomer was designated 4SE. Monomers bearing one cholesterol modification and either four or three sticky ends were named Chol-4SE and Chol-3SE, respectively. When cholesterol was positioned directly adjacent to a sticky end, the variant was referred to as Chol’-4SE. The monomer containing two cholesterol modifications and three sticky ends was denoted 2Chol’-3SE. To visualize the condensates, one DNA strand in each monomer was labeled with a Cy3 fluorophore (Table S1). Confocal laser scanning microscopy (CLSM) images revealed the formation of spherical condensate droplets with an average diameter of 3-4 μm (Fig. S1, 2).

To examine the liquid-like behaviors of the designed TDC droplets, we performed fluorescence recovery after photobleaching (FRAP) on Cy3-labeled droplets. All droplets displayed fluorescence recovery within the photobleached regions (white circles), confirming dynamic molecular exchange within the condensates (Fig. 1b and 1c). The recovery efficiencies followed the order 4SE (95%) > Chol’-4SE (75%) > Chol-3SE (65%) > 2Chol’-3SE (52%) > Chol-4SE (46%). The corresponding half-recovery times (t_₁/₂_, min) were 3.52 min for 4SE, 4.06 min for Chol’-4SE, 6.26 min for Chol-3SE, 7.13 min for 2Chol’-3SE and 13.47 min for Chol-4SE. Unmodified 4SE droplets exhibited the fastest and most complete fluorescence recovery, indicating the highest internal fluidity, whereas cholesterol-functionalized droplets displayed reduced recovery and restricted molecular mobility. The lower recovery of 2Chol’-3SE and Chol-4SE further suggests that cholesterol incorporation, cholesterol valency and monomer architecture collectively regulate internal droplet dynamics.

Notably, Chol’-4SE droplets recovered faster than Chol-3SE droplets, despite containing more sticky ends. This comparison suggests that not only the number of sticky ends, but also their spatial arrangement relative to cholesterol, controls condensate mobility. We hypothesize that removal of the sticky end adjacent to cholesterol in Chol-3SE enhances local cholesterol-cholesterol interactions or hydrophobic clustering, thereby restricting molecular exchange within the droplets^26^. By contrast, the adjacent sticky-end design in Chol’-4SE may help disperse cholesterol within the DNA network, maintaining higher internal fluidity. This observation provides a direct rationale for further examining cholesterol organization within the condensates.

To independently assess condensate fluidity, we analyzed droplet fusion dynamics. TDC droplets readily fuse into larger droplets and the relaxation of fused droplets from ellipsoidal to spherical shapes was quantified by tracking the time-dependent change in aspect ratio. Because shape relaxation after droplet fusion depends on droplet size and the inverse capillary velocity (η/γ, where η is the apparent viscosity and γ is the interfacial tension)^2, 27^, we extracted η/γ values from the relationship between relaxation time and characteristic length (Fig. 1d and S3). The η/γ values increased in the order 4SE (6.94) < Chol’-4SE (8.02) < Chol-3SE (13.92) < 2Chol’-3SE (17.43) < Chol-4SE (26.68), consistent with the FRAP results. Lower η/γ values correspond to faster relaxation and higher apparent fluidity, whereas higher η/γ values indicate slower relaxation and more viscous or less dynamic droplets. The fluidic and fusion properties of the constructed droplets are summarized in Table S2. These results indicate that molecular programming of sticky-end valency, cholesterol valency and cholesterol positioning enables systematic tuning of DNA condensate material properties, including molecular mobility and fusion dynamics. These differences in liquid-like behavior and internal composition provide the material basis for the programmed membrane-interaction states examined below.

### Cholesterol organization regulates DNA condensate mobility

Based on our earlier observations of droplet fluidity, we hypothesized that the spatial distribution of cholesterol within TDC droplets depends on both its valency and positional arrangement. To examine this, we stained the droplets with CellMask™, an amphipathic membrane dye that partitions into hydrophobic regions within the condensates (Fig 2a and Fig. S4). Interestingly, although Chol’-4SE and Chol-3SE contain the same number of cholesterol modification (with differences only in the modification position), Chol’-4SE exhibited a more homogeneous internal distribution and higher molecular mobility than Chol-3SE, suggesting that retaining a sticky end adjacent to cholesterol helps integrate cholesterol-bearing motifs into the DNA interaction network and reduces local hydrophobic clustering. By contrast, removal of the adjacent sticky end in Chol-3SE may promote cholesterol-cholesterol association or local hydrophobic enrichment, thereby restricting molecular exchange within the condensate. This interpretation is consistent with the FRAP results, in which Chol’-4SE recovered faster than Chol-3SE despite containing more sticky ends.

**Fig. 2.**
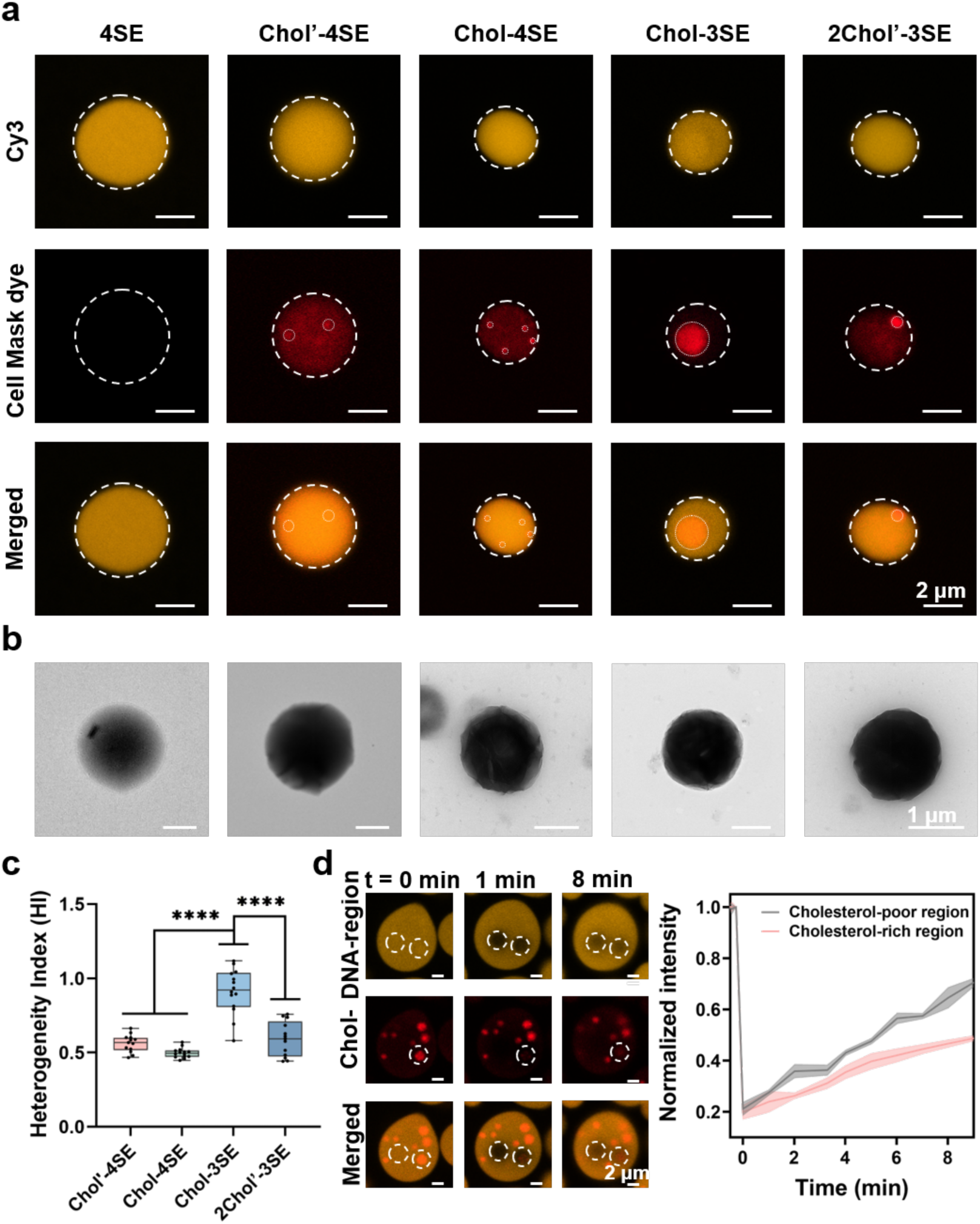
Cholesterol distribution and morphology of TDC droplets. (a) Representative CLSM images displaying cholesterol distribution within different TDC droplets. Cy3-labeled droplets are shown in yellow, and cholesterol is visualized with a lipid-specific fluorescent dye (red). Dashed outlines indicate droplet boundaries. Scale bars: 2 μm. (b) TEM images of corresponding TDC droplets showing their spherical morphology and the effect of cholesterol incorporation. Scale bars: 1 μm. (c) Quantification of cholesterol distribution using heterogeneity index (HI) (see Methods, N > 20). ****, P < 0.0001, one-way ANOVA. (d) Regional FRAP analysis within Chol-3SE droplets. Cholesterol-poor and cholesterol-rich regions were selected within the same droplets, and Cy3 fluorescence recovery was monitored after photobleaching. Recovery curves show slower molecular exchange in cholesterol-rich regions than in cholesterol-poor regions. Scale bars: 2 μm.

Morphological analysis of TDC droplets using transmission electron microscopy (TEM) further displayed all droplets maintained generally spherical morphologies, consistent with confocal microscopy observations (Fig. 2b). Notably, the high local concentration of DNA within the phase-separated droplets allowed clear visualization without additional staining. Cholesterol-modified droplets exhibited stronger electron contrast than unmodified droplets, possibly reflecting differences in local density, drying morphology or hydrophobic organization after cholesterol incorporation. Combining fluorescence imaging with these observations reveals that DNA sequence design and cholesterol positioning regulate internal condensate organization.

Moreover, we quantified the spatial heterogeneity of cholesterol-associated dye signals within individual droplets using a heterogeneity index (HI), calculated from the dye signal within droplet regions defined by the Cy3 channel (see Methods). As shown in Fig. 2a and 2c, Chol-3SE droplets, in which the sticky ends adjacent to cholesterol was removed, displayed clear cholesterol clustering, as evidenced by large red fluorescent aggregates within the droplets and the highest HI. In contrast, Chol’-4SE, retaining the adjacent sticky end, displayed a more dispersed cholesterol distribution and a lower HI value, although minor local clustering was still observed. Chol-4SE droplets exhibited relatively low HI, suggesting less heterogeneous hydrophobic organization despite their reduced droplet fluidity. For 2Chol’-3SE, which contains both positional configurations of cholesterol, staining revealed a mixed pattern of local aggregates and dispersed signals, corresponding to an intermediate HI. These results support our hypothesis that positional arrangement governs cholesterol organization within droplets.

To directly assess how local cholesterol enrichment affects molecular mobility, we performed regional FRAP analysis within Chol-3SE droplets that exhibited heterogeneous cholesterol distribution. Cholesterol-rich and cholesterol-poor regions were selected within the same droplets and monitored after photobleaching. As shown in Fig. 2d, fluorescence recovery in cholesterol-rich regions was much slower than in cholesterol-poor regions, indicating that local cholesterol enrichment is associated with restricted molecular exchange and reduced local fluidity. These results demonstrate that molecular programming of cholesterol position and sticky-end arrangement enables the tuning of both global droplet fluidity and local internal organization, providing a material basis for the distinct membrane-interaction states examined below.

### Programmed membrane wetting of DNA condensate droplets at living cell membranes

Having established that sticky-end valency and cholesterol organization tune the material properties of TDC droplets, we explored whether these parameters regulate condensate interactions with living cell membranes. Condensate-membrane wetting has been quantitatively described in reconstituted membrane systems, but how programmable condensates interact with active plasma membranes remains less understood. We therefore used RAW 264.7 macrophages as a living cell interface with active membrane dynamics and robust endocytic activity^28^, and monitored the interaction of different TDC droplets with CellMask™-labeled plasma membranes over two hours.

Unmodified 4SE droplets revealed only transient or sporadic contacts with the cell surface and largely retained their spherical morphology, indicating a weak-contact state (Fig. 3a, b and Video S1). In contrast, cholesterol-modified droplets exhibited markedly enhanced membrane engagement. Chol’-4SE droplets rapidly spread along the plasma membrane and showed clear deformation at the contact region, consistent with a wetting-dominated interaction state (Fig. 3b and Video S2). Chol-3SE droplets also interacted with the membrane, but with slower and less extensive wetting. By comparison, Chol-4SE and 2Chol’-3SE droplets exhibited persistent surface association with limited spreading, suggesting a surface-anchoring state rather than full wetting (Fig. 3b and Videos S3-S5). While these observations corroborate the trends in droplet fluidity, they further reveal that fluidity alone is insufficient to govern membrane wetting. Instead, the spatial organization of cholesterol within the condensates emerges as a key determinant of wetting and anchoring states. Among the cholesterol-modified formulations, Chol’-4SE displayed the highest fluidity and the most rapid wetting kinetics. However, droplets with reduced fluidity or stronger cholesterol clustering showed persistent surface anchoring rather than extensive spreading. These results suggest that membrane wetting is governed by the combined effects of cholesterol-mediated membrane affinity, droplet fluidity and cholesterol internal organization. Unlike reconstituted membrane systems, where wetting transitions are often induced by changing buffer conditions, temperature or membrane composition^29^, cholesterol-programmed TDC droplets engaged macrophage plasma membranes under constant physiological conditions. This behavior reflects the interplay between material properties and active membrane dynamics in living cells^30^.

**Fig. 3.**
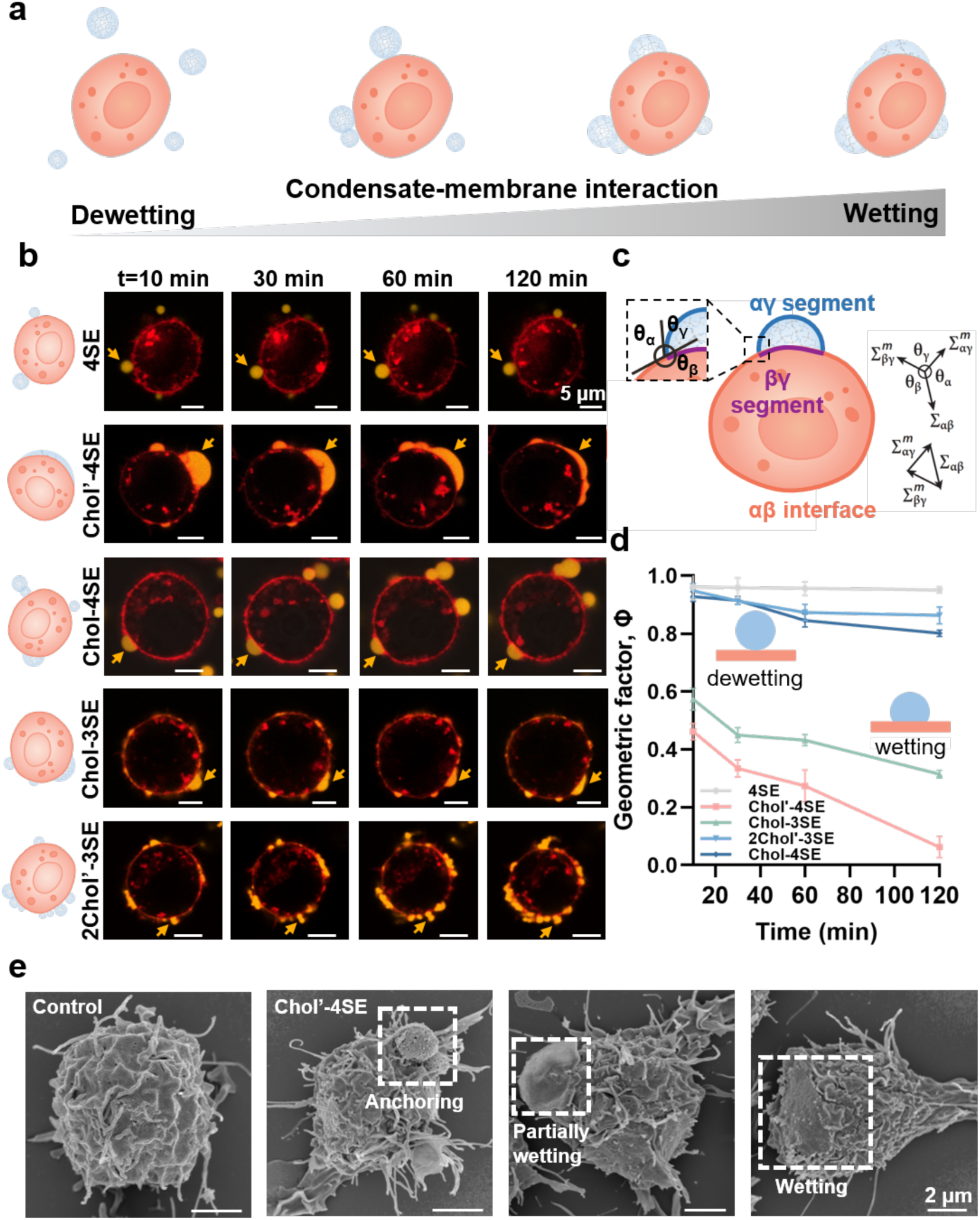
Programmed wetting and anchoring of TDC droplets at macrophage plasma membranes. (a) Schematic illustration of distinct condensate-membrane interaction states, ranging from weak contact to membrane wetting and surface anchoring. (b) Time-lapse CLSM images of Cy3-labeled TDC droplets (orange) interacting with CellMask™-stained RAW 264.7 plasma membranes (red). Yellow arrows indicate representative droplets at the membrane interface. Scale bars, 5 μm. (c) Schematic of the Young-Neumann force balance used for apparent contact-angle analysis. The αγ segment, αβ interface and βγ segment are indicated in blue, orange and purple, respectively, and define the apparent contact angles θ_α_, θ_β_ and θ_γ_ at the three-phase contact line. (d) Time-dependent changes in the geometric factor Φ for different TDC droplets. Decreasing Φ indicates increased apparent wetting. Data represent mean ± s.d. from three independent experiments. (e) Representative SEM images showing sequential membrane-engagement states of Chol’-4SE droplets on RAW 264.7 cells. Scale bars, 2 μm.

To quantify these interaction states, we adapted contact-angle analysis from established condensate-membrane wetting models^14, 16^. In a partially wetted configuration, the condensate interface and the two membrane-associated segments meet at a three-phase contact line, where the apparent contact angles reflect the balance of interfacial tensions described by Young-Neumann geometry (Fig. 3c and Methods). For droplets remaining at the cell surface, the apparent angles were measured from confocal images by tracing the local tangent directions of the droplet contour and the plasma membrane at the contact region. Because living cell membranes are active, heterogeneous and mechanically coupled to the cytoskeleton, we used the derived parameters as apparent geometric descriptors of wetting morphology rather than absolute intrinsic material constants.

Using this framework, we quantified the time-dependent apparent interfacial contact angle to compare condensate-membrane engagement and wetting (Fig. S5 and S6). Because living plasma membranes are actively regulated by cytoskeletal forces and possess complex local curvatures, we emphasize that these parameters serve as operational geometric indices of the wetting morphology rather than absolute thermodynamic material constants. Chol’-4SE droplets showed the largest decrease in contact angle and the highest apparent adhesion affinity, followed by Chol-3SE, whereas Chol-4SE and 2Chol’-3SE displayed more limited interfacial deformation. Notably, 2Chol’-3SE droplets, despite their increased cholesterol content, preferentially formed a persistent surface-anchoring state instead of undergoing extensive membrane wetting (Fig. 3b, fifth row). These results support the conclusion that wetting is not simply enhanced by increasing cholesterol valency, but is governed by the combined effects of droplet fluidity and internal cholesterol organization.

We further introduced a geometric factor (Φ), as an operational descriptor of condensate-membrane wetting. In this analysis, Φ values approaching 1 correspond to weak contact or dewetting, whereas decreasing Φ values indicate enhanced apparent wetting. As shown in Fig. 3d, unmodified 4SE droplets remained close to the dewetting regime, while Chol’-4SE droplets underwent the most repid decrease in Φ over time. Chol-3SE occupied an intermediate wetting regime, whereas Chol-4SE and 2Chol’-3SE stayed in a weakly deforming or anchoring-dominated state. Importantly, these membrane-interaction states mirrored the material properties of the droplets. Chol’-4SE droplets, which combined relatively high fluidity with dispersed cholesterol organization, exhibited the strongest membrane wetting. Conversely, droplets with lower fluidity or enhanced cholesterol clustering spread inefficiently and tended toward weak contact or surface anchoring (Fig. 1c, 2 and 3d).

Scanning electron microscopy (SEM) further supported the physical engagement between TDC droplets and living cell membranes. After incubation with Chol’-4SE droplets, micron-sized condensates were observed on the macrophage surface, with filopodia-like membrane protrusions extending toward and around the droplets (Fig. 3e). These structures are consistent with active membrane participation during droplet capture and internalization, and support the view that programmed condensate wetting is coupled to cellular membrane machineries.

Recent GUV-based studies have displayed that membrane lipid packing and cholesterol content can regulate condensate wetting affinity^16^. Increased cholesterol can alter interfacial water organization and membrane dipole potential^31^, which may facilitate the selective condensate association with organelles that differ in cholesterol content^32^. Moreover, membrane cholesterol content is known to modulate membrane fluidity^33^. We examined whether TDC interactions with living cell membranes are also sensitive to native membrane composition. Chol-3SE droplets were further assessed in RAW 264.7, HeLa and 4T1 cells, which represent different native membrane environments with reported differences in membrane lipid composition, cholesterol abundance and lateral organization^34, 35^. After 2 h of incubation, Chol-3SE droplets showed clear wetting on RAW 264.7 cells but only occasional anchoring was observed on HeLa and 4T1 cells (Fig. S7). These observations suggest that cell type specific membrane composition and organization influence condensate wetting and anchoring at living cell interfaces, consistent with membrane composition dependent wetting behavior observed in reconstituted systems^16^.

### Native membrane composition modulates condensate wetting and anchoring

Having observed condensate type dependent differences in mammalian cells, we next asked whether native membrane composition could modulate condensate wetting and anchoring in a broader context. Plant plasma membranes provide a useful comparison because they contain sterol-rich but compositionally different lipid environments relative to mammalian membranes^36^. We therefore examined TDC interactions with protoplasts derived from *Nicotiana benthamiana* (*N. benthamiana*) callus tissue, in which the cell wall is removed and the plasma membrane is directly accessible.

Three representative TDC droplets were selected for this analysis: unmodified 4SE droplets, which showed weak interactions with macrophage membranes, and two cholesterol-modified droplets, Chol’-4SE and Chol-3SE, which displayed stronger membrane engagement in mammalian cells. Protoplasts were labeled with fluorescein diacetate (FDA), and Cy3-labeled droplets were monitored over time. Consistent with the mammalian cell observations, 4SE droplets exhibited minimal contact with callus-derived protoplasts, whereas Chol’-4SE and Chol-3SE droplets exhibited clear membrane interactions (Fig. 4a, b). However, protoplast-droplet interactions developed more slowly and displayed less extensive wetting within the first two hours. Images of extended incubation time revealed a progressive decrease in the geometric factor Φ after 6 h, indicating increased apparent wetting and membrane engagement (Fig. 4c). Chol’-4SE droplets were frequently internalized after 12 h, whereas Chol-3SE droplets were internalized more slowly, typically after 24 h. These results suggest that cholesterol-modified TDC droplets can engage plant protoplast membranes, but with slower wetting-to-uptake kinetics than those observed in macrophages. There is significant diversity in plasma-membrane lipid composition among plant species^37^ and among different tissues of the same species^38^. This prompted us to compare protoplasts derived from *N. benthamiana* callus and leaves to examine whether native lipidomic differences influence condensate-membrane interactions. Under identical incubation conditions, Chol’-4SE droplets interacted strongly with callus-derived protoplasts but showed little apparent interaction with leaf-derived protoplasts (Fig. 4d). Lipidomic analysis by mass spectrometry identified 1,239 lipid metabolites in positive ion mode and 534 lipid metabolites in negative ion mode, revealing clear compositional differences between callus- and leaf-derived protoplasts. Callus-derived protoplasts showed higher relative abundance of several short-chain and saturated lipid species, including diacylglycerols (DG, 6:0/9:0), phosphatidylcholines (PC), lysophosphatidylethanolamines (LPE) and phosphatidylglycerols (PG). In contrast, leaf-derived protoplasts were enriched in more unsaturated and long-chain lipid species, including diacylglycerols (DG, 22:4/12:2), monogalactosyldiacylglycerols (MGDG), phosphatidylinositols (PI) and phosphatidylethanolamines (PE). These lipidomic differences provide a physical basis for interpreting the differential condensate-membrane behaviors observed in callus- and leaf-derived protoplasts. Lipid chain saturation, chain length and sterol compatibility can influence membrane packing, local order and hydrophobic insertion^39–42^. Specifically, short-chain lipids, such as DG, could increase local membrane fluidity, facilitate cholesterol insertion and promote faster integration of cholesterol into the membrane^41^. Saturated/monounsaturated lipids, such as PC, can form more ordered lipid raft domains, favoring cholesterol anchoring^39^. We therefore propose that callus-derived protoplasts with higher abundance of several short-chain and saturated lipid species provide a more favorable lipid environment for cholesterol-modified DNA condensates to anchor and subsequent wetting. In contrast, leaf-derived protoplasts possessing more unsaturated and long chain lipid species may reduce stable cholesterol-mediated engagement. Our results suggest that native lipid composition is an important modulator of DNA-condensate wetting and anchoring at living cell membranes.

**Fig. 4.**
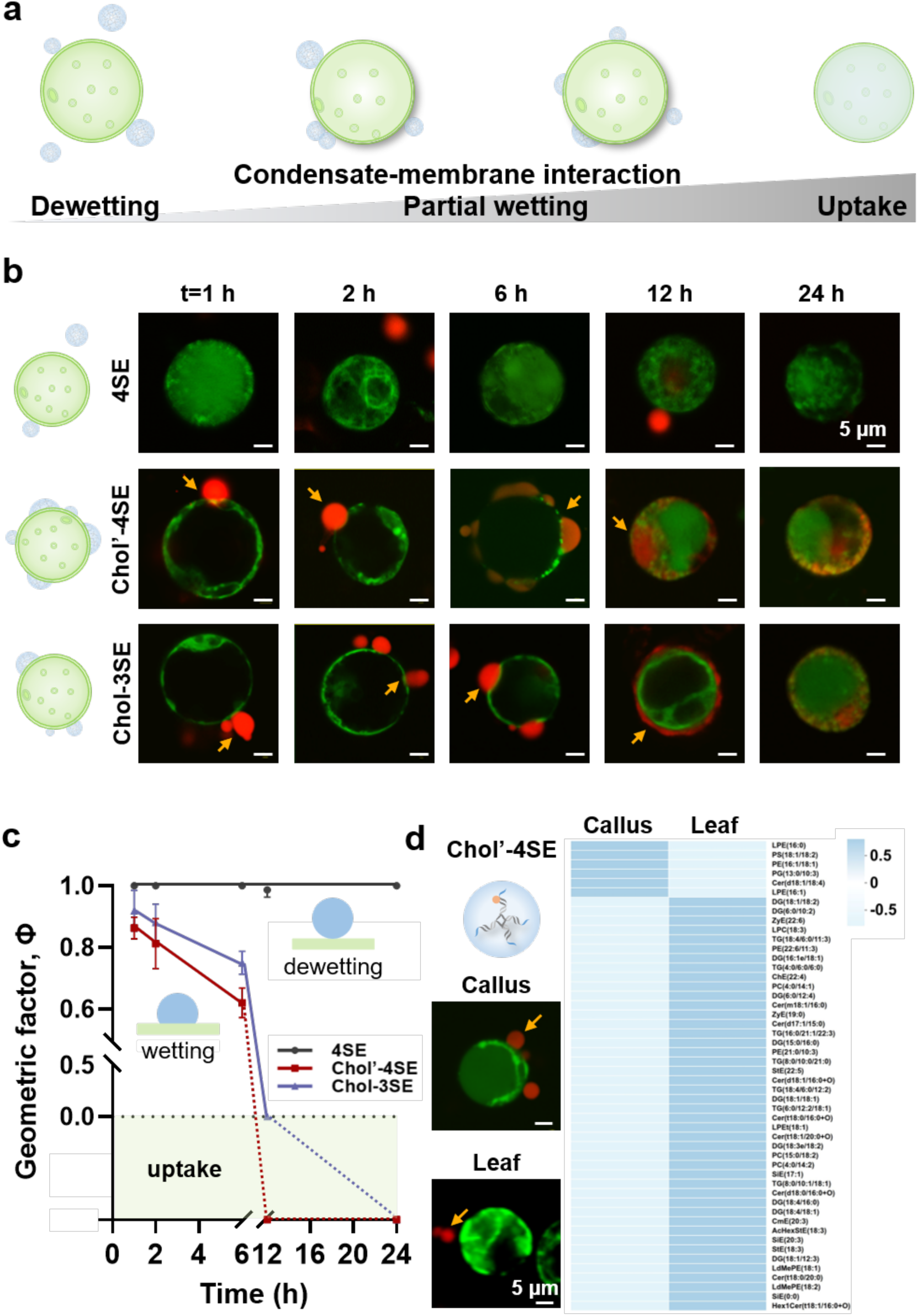
Native membrane composition modulates TDC wetting and anchoring in plant protoplasts. (a) Schematic illustration of condensate-protoplast interaction states, ranging from weak contact to partial wetting and uptake. (b) Time-lapse CLSM images of Cy3-labeled TDC droplets interacting with FDA-stained *N. benthamiana* callus-derived protoplasts. Red, TDC droplets; green, protoplasts. Yellow arrows indicate representative condensate-membrane contacts. Scale bars, 5 μm. (c) Time-dependent changes in the apparent geometric factor Φ for different TDC droplets interacting with callus-derived protoplasts. Decreasing Φ indicates increased apparent membrane engagement; uptake events were analyzed separately from surface wetting states. Data represent mean ± s.d. from three independent experiments. (d) Representative CLSM images showing differential interactions of Chol’-4SE droplets with callus- and leaf-derived protoplasts under identical incubation conditions, together with lipidomic comparison of the two protoplast sources by mass spectrometry. Scale bars, 5 μm.

Across mammalian cell and plant protoplast, TDC membrane engagement is controlled by both condensate properties and native membrane composition. Cholesterol-modified droplets with relatively dispersed internal cholesterol organization and higher fluidity favor rapid wetting, whereas increased cholesterol valency or stronger local cholesterol clustering biases droplets toward surface anchoring. In plant protoplasts, membrane composition further modulates these interaction states and the kinetics of internalization. Notably, membrane invagination was observed adjacent to Chol-3SE droplets during protoplast engagement at 6 h (Fig. 4b), suggesting that condensate wetting can couple to membrane deformation in this native membrane context. These findings extend the programmable wetting behavior of DNA condensates from macrophage plasma membranes to compositionally different living cell membranes.

### Wetting competence distinguishes surface anchoring from cellular internalization

Condensate droplets can wet and deform lipid membranes and undergo internalization via endocytosis, a process driven by condensate-membrane interactions^43, 44^. We explored whether these programmed interfacial states influence cellular uptake and internalization of TDC droplets. RAW 264.7 cells were incubated overnight with Cy3-labeled TDC droplets, and living cell membranes were stained using CellMask™ dye. CLSM images revealed efficient internalization of all droplet types, with cholesterol-modified droplets displaying higher cellular accumulation than unmodified 4SE droplets (Fig. 5a and Fig. S8). Three-dimensional z-stack imaging further confirmed the intracellular localization of Chol’-4SE droplets, supporting their uptake as micron-scale condensates rather than simple surface adsorption (Fig. 5b). Quantitative image analysis (Fig. 5c) revealed that cellular uptake efficiency correlated with the membrane-interaction states defined above (Fig. 3d). Chol’-4SE and Chol-3SE droplets, which displayed extensive membrane wetting, showed the highest uptake efficiencies. In contrast, 2Chol’-3SE droplets, despite containing a higher cholesterol valency, preferentially formed surface-anchoring states and did not further enhance uptake. (Fig. 3b and Fig. 5c). Chol-4SE droplets, which showed lower fluidity and weaker membrane deformation, and unmodified 4SE droplets, which displayed weak membrane contact, exhibited lower uptake. These results indicate that efficient internalization requires not only membrane association, but also an appropriate wetting or deformable anchoring state at the plasma membrane.

**Fig. 5.**
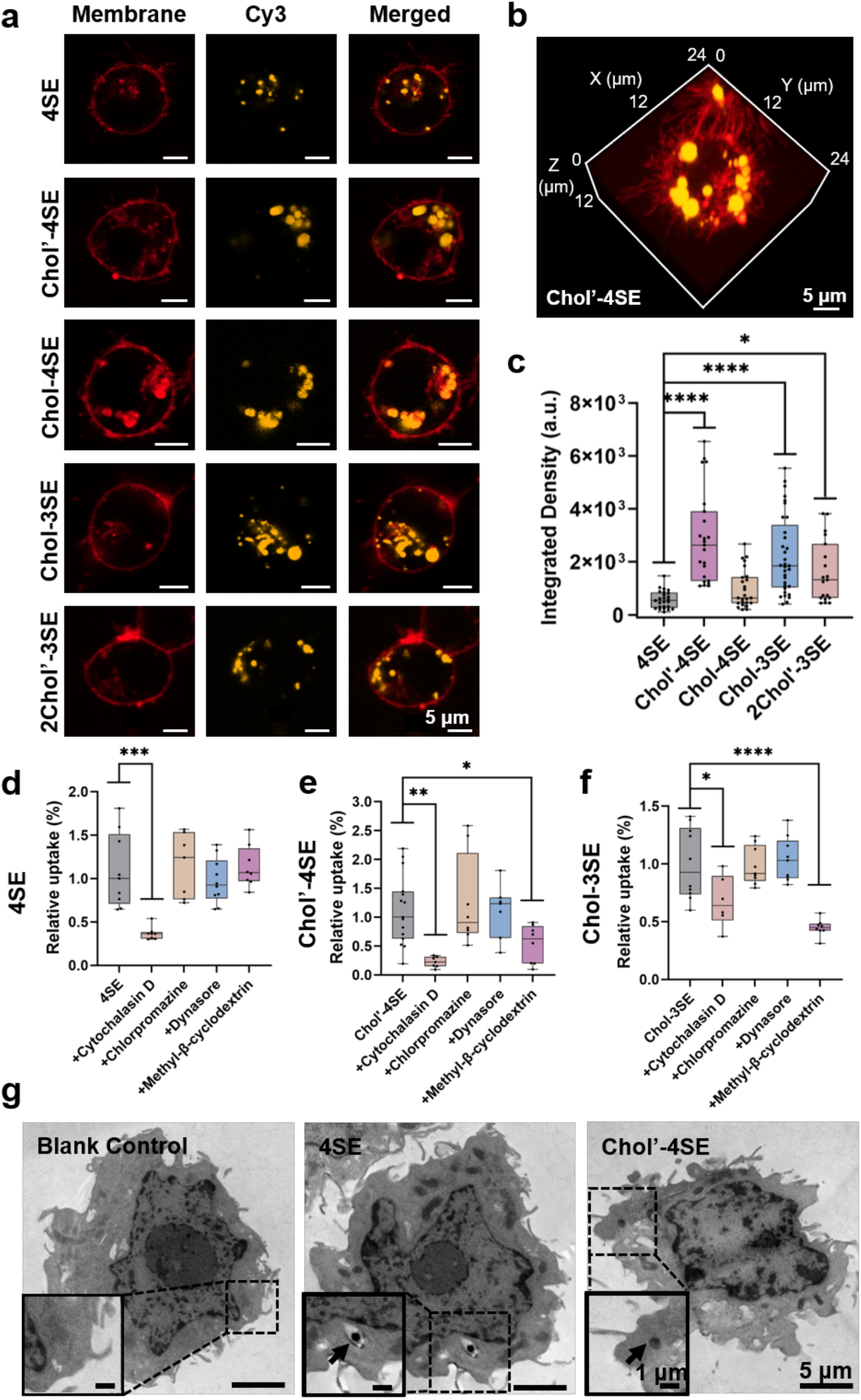
Wetting competence predicts the transition from surface association to cellular internalization. (a) Representative confocal images showing uptake of different Cy3-labeled TDC droplets (orange) by RAW 264.7 cells. Plasma membranes were labeled with CellMask™ (red). Scale bars: 5 μm. (b) Three-dimensional z-stack reconstruction confirming intracellular localization of Chol’-4SE droplets. Scale bar: 5 μm. (c) Quantitative analysis of intracellular Cy3 fluorescence intensity for different TDC types (N > 20 cells per group). ****, P < 0.0001; *, P < 0.05, one-way ANOVA. (d-f) Quantification of relative fluorescence intensity for different TDC droplets after pretreatment with four different inhibitors: Cytochalasin D, chlorpromazine, dynasore, and methyl-β-cyclodextrin (M-β-CD). Data represent mean ± s.d. from three independent experiments. Statistical significance was determined by one-way ANOVA: ****P < 0.0001, ***P < 0.001, **P < 0.01, *P < 0.05. (g) TEM images of cells incubated with Chol’-4SE or 4SE droplets for 12h. Enlarged images highlight intracellular or membrane-associated droplets (black arrows). Scale bars: 5 μm (overview) and 1 μm (enlarged).

To examine the cellular machineries involved in droplet internalization, we pretreated cells with pharmacological inhibitors before droplet incubation (see details in Methods). Cytochalasin D was used to disrupt actin polymerization, chlorpromazine to inhibit clathrin-mediated endocytosis, dynasore to inhibit dynamin-dependent uptake, and methyl-β-cyclodextrin (M-β-CD) to deplete membrane cholesterol and perturb lipid rafts and caveolae^45^. Quantitative fluorescence analysis showed that chlorpromazine and dynasore had limited effects on the uptake of most TDC droplets (Fig. S9). By contrast, Cytochalasin D significantly reduced cellular uptake across all droplets, indicating a major contribution of actin-dependent internalization (Fig. 5d-f). For unmodified 4SE droplets, Cytochalasin D was the only inhibitor that markedly reduced uptake (Fig. 5d), suggesting actin-dependent engulfment-like entry mode. For cholesterol-modified droplets, the inhibitory effect of Cytochalasin D also supports our SEM observations exhibiting filamentous membrane protrusions engaging Chol’-4SE droplets at the cell surface (Fig. 3e), confirming the involvement of actin remodeling during wetting-associated internalization. Meanwhile, the sensitivity of cholesterol-modified droplets to M-β-CD indicates that their uptake is responsive to membrane cholesterol depletion, potentially through cholesterol-rich plasma membrane microdomains such as caveolae or lipid rafts^46^. Notably, although Chol’-4SE and Chol-3SE exhibited comparable uptake efficiencies, they showed different inhibitor sensitivities, with Cytochalasin D more strongly reducing Chol’-4SE uptake and M-β-CD producing stronger inhibition of Chol-3SE uptake (Fig. 5e, f). We attribute this difference to differential cholesterol organization within the droplets. Chol-3SE droplets exhibit stronger local cholesterol clustering (Fig. 2a), which may favor more localized interactions with cholesterol-enriched membrane regions during internalization.

TEM images further supported the intracellular localization of TDC droplets after 12 h incubation (Fig. 5g). Unmodified 4SE droplets were observed within vesicle-like compartments, whereas Chol’-4SE droplets appeared closely associated with membrane protrusions or filamentous structures^25^, consistent with their stronger membrane engagement. Together, these results suggest that TDC droplet internalization is not a purely passive uptake process, but is associated with active membrane remodeling. Molecularly programmed apparent wetting and anchoring geometries shape the initial condensate-membrane contact state and influence differential actin-dependent cellular entry. This supports a mechanobiological model in which the physical state of the condensate interface biases membrane engagement, cytoskeletal recruitment and internalization behavior. Rather than following a single canonical endocytic pathway, TDC droplets appear to enter cells through interaction state dependent modes, with wetting-associated droplets favoring actin-dependent engulfment and cholesterol-clustered droplets showing stronger cholesterol-sensitive uptake. These findings provide mechanistic insight into how condensate composition and interfacial state shape cellular entry, offering design principles for programmable condensate-based delivery systems.

### Condensate physical integrity governs functional cargo delivery

Building on our finding that programmed membrane wetting promotes cellular internalization, we then asked whether TDC droplets could harness this entry behavior to deliver functional biomolecules. We selected Chol’-4SE droplets as the carrier due to their strong membrane wetting and excellent cellular uptake efficiency. The droplets were loaded with three functional biomolecules, CpG ODNs, luciferase mRNA and streptavidin (SA) protein, *via* the LLPS process through rational sequence design and functional modification. Specifically, CpG ODNs were incorporated into DNA monomers to enrich immunostimulatory sequences within condensates. The polyT30 domain was introduced to hybridize with the polyA tail of mRNA and biotinylated DNA monomers were used to recruit SA through biotin-streptavidin binding (Fig. 6a).

**Fig. 6.**
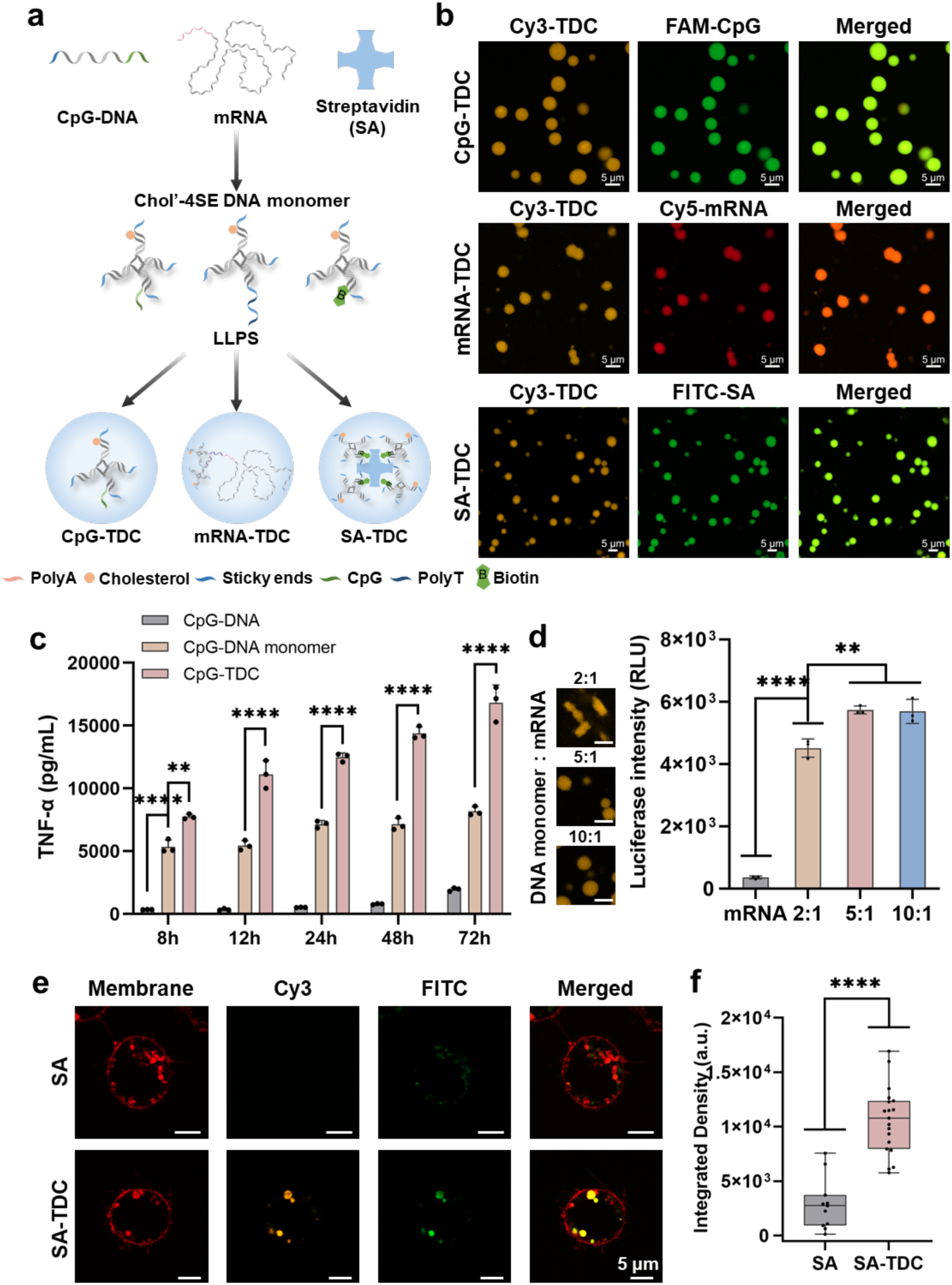
Wetting-enabled enrichment and intracellular delivery of functional biomolecules by TDC droplets. (a) Schematic illustration of cargo incorporation into Chol’-4SE droplets. CpG ODNs were incorporated through DNA sequence design, mRNA was recruited through polyA/polyT hybridization, and SA protein was incorporated through biotin-streptavidin binding. (b) Fluorescence imaging showing co-localization of Cy3-labeled TDC droplets with FAM-labeled CpG ODNs, Cy5-labeled mRNA or FITC-labeled SA. Scale bar: 5 μm. (c) TNF-α secretion from RAW 264.7 cells treated with free CpG ODNs, CpG-conjugated DNA monomers or CpG-loaded TDC droplets at a final CpG concentration of 100 nM. Data analyzed by one-way ANOVA; **, P < 0.01; ****, P < 0.0001. (d) Luciferase activity after delivery of luciferase mRNA using TDC droplets formed at different DNA monomer-to-mRNA ratios (n = 3). Representative images (left) show the morphology of mRNA-loaded droplets. Scale bars: 5 μm. (e) CLSM images of RAW 264.7 cells incubated with free SA or SA-loaded TDC droplets. Scale bar: 5 μm. (f) Quantification of intracellular FITC-SA fluorescence intensity (N = 10). Data represent mean ± s.d.; statistical significance was determined by one-way ANOVA: ****P < 0.0001, ***P < 0.001, **P < 0.01, *P < 0.05.

Fluorescence imaging confirmed efficient co-localization of FAM-labeled CpG ODNs, Cy5-labeled mRNA and FITC-labeled SA with Cy3-labeled Chol’-4SE droplets (Fig. 6b). These results demonstrate that DNA condensate droplets can enrich chemically and structurally different biomolecules without disrupting condensate formation. For mRNA loading, the large size of the ∼2000 nt transcript affected droplet morphology at low DNA monomer-to-mRNA ratios. Condensates formed at a 2:1 ratio showed irregular morphologies, whereas ratios of 5:1 and 10:1 produced more spherical droplets (Fig. 6d and Fig. S10). Gel electrophoresis further confirmed efficient mRNA incorporation into the droplets (Fig. S11), indicating that the modular DNA design enables cargo enrichment while preserving the droplet state when the monomer-to-cargo ratio is appropriately controlled. This morphology dependence suggests that excessive mRNA incorporation can perturb the condensate architecture and compromise the interfacial properties required for productive membrane engagement. Consistently, droplets formed at 5:1 and 10:1 ratios generated higher luciferase expression than irregular condensates formed at the 2:1 ratio. These results indicate that functional mRNA delivery is not governed solely by cargo loading, but also by the physical state of the condensate, with droplet morphology and interfacial integrity shaping the intracellular fate of macromolecular cargo.

CpG ODNs are recognized by Toll-like receptor 9 in immune cells, triggering innate and adaptive immune responses, including the production of pro-inflammatory cytokines such as tumor necrosis factor-alpha (TNF-α)^47, 48^. However, free CpG ODNs are susceptible to nuclease degradation and exhibit poor cellular uptake under physiological conditions. We first evaluated CpG ODN delivery in RAW 264.7 macrophages by measuring TNF-α secretion. As shown in Fig. 6c, after 8 h of incubation, free CpG ODNs elicited minimal immune stimulation, whereas CpG-TDC droplets induced approximately 1.5-fold higher TNF-α secretion than CpG-conjugated monomers, indicating enhanced immunostimulatory activity of the condensate formulation. TNF-α production continued to increase from 12 to 72 h in cells treated with CpG-TDC droplets compared with CpG-conjugated monomers, suggesting that condensate loading improved CpG protection and prolonged cellular stimulation under comparable cellular uptake conditions (Fig. S8 and S12). Unmodified 4SE TDC droplets also prolonged CpG-induced TNF-α production compared with CpG-conjugated monomers, further supporting the contribution of the condensate state to sustained immunostimulatory output (Fig. S13).

We next assessed delivery of luciferase mRNA as a functional nucleic acid cargo. Luciferase mRNA (∼2000 nt) was selected as a model cargo because the delivery of large and structurally fragile nucleic acids remains challenging owing to their susceptibility to degradation and limited cellular uptake^49^. Compared with free mRNA, mRNA-loaded TDC droplets produced significantly higher luciferase activity after 48 h, confirming that the droplets enabled intracellular mRNA delivery while preserving translational activity (Fig. 6d). The dependence of luciferase expression on droplet morphology, described above, further supports the importance of condensate architecture and interfacial integrity for functional mRNA delivery. In plant protoplasts, mRNA-loaded TDC droplets also enhanced luciferase expression compared with free mRNA (Fig. S14), supporting the potential generality of this delivery strategy in compositionally different cellular systems.

Finally, we evaluated protein internalization using SA as a model cargo. FITC-labeled SA was efficiently recruited into droplets through biotinylated DNA monomers and delivered to RAW 264.7 cells. Compared with free SA, SA-loaded TDC droplets produced stronger intracellular FITC signals with punctate distribution (Fig. 6e and Fig. S15). Quantitative fluorescence analysis confirmed approximately fourfold higher cellular internalization of SA when delivered by TDC droplets (Fig. 6f). These data demonstrate that TDC droplets can enhance intracellular accumulation of protein cargoes, although further studies will be required to define the subcellular release and functional activity of delivered proteins.

By integrating programmable cargo enrichment with wetting-enabled cellular uptake, TDC droplets enabled the intracellular delivery of three types of biomolecules, including oligonucleotides, mRNA and proteins. These results indicate that TDC droplets can preserve functional cargo responses and, in the case of CpG ODNs, support prolonged immunostimulatory activity, likely through enhanced cargo protection and sustained cellular access. This multifunctionality positions TDC droplets as a promising programmable delivery platform for applications requiring modular cargo loading and prolonged biological activity.

## Discussion

This work engineers DNA-programmed condensate droplets as a tunable material platform for controlling interfacial fate at living cell membranes. By varying sticky-end valency together with cholesterol valency and positioning, we programmed different condensate properties, including molecular mobility, fusion dynamics and cholesterol organization. These programmed material states generated three distinct membrane-interaction behaviors, weak contact, rapid wetting and persistent surface anchoring. The results define how sequence-defined molecular architecture controls living membrane engagement and subsequent cellular internalization.

A crucial finding is that membrane adhesion does not necessarily progress towards productive cellular entry. Cholesterol modification enhanced membrane association, but increasing cholesterol valency did not monotonically increase wetting or uptake. Droplets with restricted molecular mobility frequently remained in persistent surface-anchoring states, whereas extensive wetting occurred when cholesterol-mediated membrane affinity was combined with sufficient condensate fluidity and interfacial deformability. These observations identify the transition from membrane adhesion to productive internalization as a critical interfacial bottleneck for deformable materials. The balance between adhesion and deformability, rather than either property alone, therefore provides a physical design rule that distinguishes surface retention from cellular entry.

Living plasma membranes are active and compositionally complex interfaces rather than passive substrates. Their lipid heterogeneity, local curvature and coupling to cytoskeletal and endocytic machineries alter the local physical environment encountered by membrane-bound condensates. Consistent with this view, wetting-associated droplets exhibited higher uptake and stronger actin sensitivity, whereas cholesterol-clustered droplets showed greater sensitivity to membrane cholesterol depletion. Moreover, the different interfacial states observed across mammalian cells and plant protoplasts indicate that condensate fate emerges from the interplay between material state and membrane context. Lipidomic differences between callus- and leaf-derived protoplasts further identify native membrane composition as an additional factor governing condensate anchoring and wetting. Wetting therefore represents an interfacial process through which molecularly encoded condensate properties influence cellular internalization behaviors.

This perspective reframes condensate-mediated biomolecule delivery. Conventional carrier optimization commonly emphasizes cargo loading, membrane affinity and receptor-mediated uptake, whereas our results identify wetting competence as an additional design parameter for soft delivery materials. TDC droplets enriched and delivered CpG ODNs, mRNA and proteins, but functional performance was not determined by cargo incorporation alone. Excessive mRNA loading perturbed droplet morphology and reduced luciferase expression, demonstrating that successful loading does not guarantee productive delivery when condensate architecture and interfacial integrity are compromised. Preserving a wetting competent material state may therefore be essential for converting membrane-bound condensates into efficiently internalized carriers.

Programmable control over condensate interfacial fate could enable soft materials designed either for persistent cell surface anchoring or for intracellular entry. Surface retained condensates may support localized signaling, immunomodulation or membrane repair, whereas wetting competent condensates may facilitate intracellular delivery of otherwise membrane impermeable cargoes. Our findings establish a structure-property-interface relationship whereby the molecular architecture of a synthetic condensate programs its physical engagement with living cells. This design framework may inform adaptive delivery systems whose interfacial behavior is matched to the physical properties of target cell membranes. Future studies combining controlled membrane perturbation, live cytoskeletal imaging and *in vivo* or *in planta* validation will further refine these design principles and enable responsive condensate materials for immunomodulation, biomolecule delivery and synthetic biology.

## Acknowledgements

This work was supported by National Natural Science Foundation of China (22377076, 32571596). We thank the support from the National Key Research and Development Program of China (2022YFA1603603), Shanghai Municipal Science and Technology Major Project, Shanghai Pilot Program for Basic Research, Shanghai Science and Technology Committee (25HC2810200) and State Key Laboratory of Synergistic Chem-Bio Synthesis, Shanghai Jiao Tong University. The language of this manuscript was polished from the original text by DeepSeek.

## Author Contribution

Huan Z. and Honglu Z. conceived the idea, designed the study and supervised the project. Z. C. conducted most of the experiments, data analysis, and wrote the manuscript. W. C., J. Y. and D. L. performed experiments of the plant cells and W. C. and J. Y. analyzed the data. Z. C., Honglu Z., C. F. M. L., and Huan Z. revised the manuscript and all authors approved the final version.

