## Supplementary Information for "DNA-Programmed Condensate-Membrane Wetting and Cellular Internalization"

### Materials and Reagents

Sodium chloride (NaCl), agarose, magnesium chloride ($MgCl_2$), 50× TAE buffer, 5× TBE buffer, 30% acrylamide solution, ammonium persulfate (APS), N, N, N', N'-tetramethylethylenediamine (TEMED), DNA markers, 6× loading buffer, Tris (hydroxymethyl) aminomethane hydrochloride buffer (Tris-HCl, pH 8.0), and fluorescein isothiocyanate-labeled streptavidin (FITC-SA) were procured from Sangon Bioengineering Technology and Services (Shanghai, China). Paraformaldehyde, glutaraldehyde, sodium cacodylate solution, cytochalasin D, chlorpromazine, dynasore, and methyl-β-cyclodextrin were obtained from Macklin Biochemical (Shanghai, China). DNA oligonucleotides were synthesized and purified by Sangon Bioengineering Technology and Services with high-performance liquid chromatography (HPLC) purification for Cy3- or cholesterol-labeled oligonucleotides. All oligonucleotides were dissolved in DNase/RNase-free water prior to use. The sequences are listed in Supplementary Table 1. Messenger RNA (mRNA) was purchased from GenScript Biotech Corporation (Nanjing, China). The RAW264.7 cells were obtained from Procell Life Science & Technology (Wuhan, China). Fetal bovine serum (FBS), antibiotics, and culture media were supplied by Thermo Fisher Scientific Inc. (Waltham, MA, USA). The tumor necrosis factor-alpha (TNF-α) ELISA kit was acquired from Beyotime Biotechnology (Shanghai, China). The luciferase reporter assay system was purchased from Promega Corporation (Madison, WI, USA). All other buffers and solutions were prepared using ultrapure water (>18.25 MΩ). All chemicals were used as received without further purification.

### Methods

#### Synthesis of DNA monomers and DNA droplets

Four-branched and five-branched DNA nanostructures were assembled by mixing the four (S1, S2, S3, S4) or five (Five-S1, Five-S2, Five-S3, Five-S4, Five-S5) oligonucleotide strands in a tube, each at a final concentration of 20 μM, dissolved in a buffer of 10 mM Tris-HCl. The mixture was heated at 95 °C for 5 minutes, then slowly denatured and annealed down to 4 °C at a rate of 1 °C/min using a PCR thermal cycler to form the four- and five-branched DNA nanostructures. For Cy3-labeled nanostructures, S1 or Five-S3 was replaced with its Cy3-conjugated counterpart (S1-Cy3 or Five-S3-Cy3, respectively). The resulting four- and five-branched DNA structures were then mixed separately in 1× PBS and incubated at 37 °C for 1 hour, at a final strand concentration of 5 μM each, to induce DNA droplet formation.

**Confocal laser scanning microscopy (CLSM) characterization**

The formation of DNA droplets was characterized using a confocal laser scanning microscope (Leica TCS SP8 STED 3X). Cy3-labeled DNA droplets were excited at 561 nm, and their emission was recorded between 570 and 620 nm. Quantitative analysis of average fluorescence intensity in the CLSM images was performed using ImageJ. Cholesterol-functionalized DNA droplets were stained using the lipophilic membrane dye CellMask™ at a 0.01× dilution. After incubating for 3 minutes, the samples were observed via CLSM.

**Fluorescence recovery after photobleaching (FRAP) analysis**

Following the formation of DNA droplets, FRAP experiments were conducted using the Leica TCS SP8 STED 3X confocal microscope. A selected region within the Cy3-labeled DNA droplet was photobleached at 561 nm with 100% laser intensity for 20 seconds to eliminate fluorescence in that target area. Post-bleaching, a time series of images was acquired at a rate of one frame per minute, monitoring fluorescence recovery for approximately 15 minutes until a plateau was reached. The mean fluorescence intensity within the bleached region was measured at each time point, allowing for construction of a fluorescence recovery curve.

**Fusion dynamics of DNA droplets**

To investigate the fusion dynamics of DNA droplets, we prepared 5 μM DNA droplets in 1× PBS buffer. The droplets were imaged at one-minute intervals using CLSM. The aspect ratio $AR(t)$ of the droplets was calculated for each image using the formula: $AR(t) = l_{long}/l_{short}$, where $l_{\mathrm{long}}$ and $l_{\mathrm{short}}$ represent the lengths of the droplet's major and minor axes, respectively. Measurements were carried out using ImageJ software. The relaxation time $\tau$ was determined by fitting the aspect ratio data to the following exponential decay expression: $AR(t) = 1+(AR(0)-1) \cdot exp(-\tau/t)$. The characteristic length scale $l$ of the fusing condensates was defined as the geometric mean: $l = [(l_{long}(t = 0) - l_{short}(t = 0)) \cdot l_{short}(t = 0)]^{1/2}$. The relationship between $\tau$ and $l$ follows the equation: $\tau = (\mu/\gamma) \cdot l + C$, where the slope of the linear fit yields the inverse capillary velocity $\eta/\gamma$. Here, $\eta$ represents the viscosity, and $\gamma$ represents the surface tension of the condensate.

**Quantification of cholesterol distribution heterogeneity within DNA condensate droplets**

Confocal images were analyzed using Image J 1.54g. Individual TDC droplets were segmented from the Cy3 channel to generate droplet region, which were then applied to the Cell Mask dye channel (indicating cholesterol). Then, the mean intensity and standard deviation of the dye signal within each droplet region were measured. The heterogeneity index (HI)is calculated as the standard deviation (StdDev) divided by the mean fluorescence intensity (Mean) of the dye signal. For each condition, at least 20 droplets from three independent experiments were analyzed.

$$\text{Heterogeneity Index (HI)} = \frac{StdDev}{Mean}$$

**Transmission electron microscopy (TEM) characterization**

The prepared DNA droplets were deposited onto carbon-coated copper grids and allowed to adsorb for approximately 5 minutes before excess solution was wicked off. To minimize salt crystal formation during drying, the grids were gently rinsed with ultrapure water and air-dried before imaging.

For cellular samples, cells were cultured in dishes and then incubated in medium containing 1 μM DNA droplets for 12 hours. After incubation, cells were washed with PBS and fixed overnight at 4 °C in a solution of 2% paraformaldehyde, 2.5% glutaraldehyde, and 0.1 M sodium cacodylate buffer. The samples were then subjected to negative staining, graded dehydration, resin embedding, ultrathin sectioning, and mounted onto copper grids for TEM analysis. TEM images were captured using a Thermo Fisher Talos L120C G2 Transmission Electron Microscope operating at an acceleration voltage of 120 kV.

**Cell culture**

RAW264.7 cells were cultured in DMEM medium containing 10% fetal bovine serum and antibiotics (100 μg/mL streptomycin and 100 μg/mL penicillin). The cells were cultured in a 5% $CO_2$ incubator at 37°C.

**Interaction between DNA droplets and living cells**

RAW264.7 cells were seeded into culture dishes compatible with confocal laser scanning microscopy and incubated overnight to allow adherence. Following three washes with 1× PBS, the cells were stained with the membrane-specific dye CellMask™ for 10 minutes, then rinsed thoroughly with PBS to remove excess dye. Preformed Cy3-labeled DNA droplets (final concentration of 1 μM) were added directly to the dish, and interaction dynamics between the DNA droplets and the cell membrane were immediately imaged using CLSM. Time-lapse imaging was typically performed after 1 hour of co-incubation to visualize and capture the interaction processes.

The protoplast cells were placed in a centrifuge tube, and 1 μM TDC droplets were added. The mixture was incubated with shaking at 37°C for 1 to 24 hours. Prior to imaging, the protoplasts were stained with the viability dye FDA for 10 minutes, and then imaged at different time points using CLSM.

**Wetting geometry and contact angles**

All wetting geometries involve three aqueous phases: the external solution $\boldsymbol{\alpha}$, the intracellular solution $\boldsymbol{\beta}$, and the droplet's internal solution $\boldsymbol{\gamma}$. These are separated by three membrane segments (see Figure 3c). The membrane interface divides into two parts at the contact line where phase $\boldsymbol{\beta}$ meets $\boldsymbol{\alpha}$. One membrane segment, labeled $\boldsymbol{\beta}_{\gamma}$, contacts the DNA droplet, while the other segment, $\boldsymbol{\alpha\gamma}$, is exposed to the external buffer.

At the three-phase contact line, the three membrane segments meet and form three distinct contact angles: $\boldsymbol{\theta}_a$, $\boldsymbol{\theta}_\beta$, and $\boldsymbol{\theta}_\gamma$, which together sum to 360° ($\theta_a + \theta_\beta + \theta_\gamma = 360°$). These contact angles correspond-via a tension triangle (Neumann's triangle)-to three interfacial tensions and are also related to the relative affinity **W**, as defined in Equation (1). The contact angles were measured following the methods described in the referenced literature [1].

$$\Sigma_{\alpha\gamma}^{m} = \Sigma + W_{\alpha\gamma} \text{ and } \Sigma_{\beta\gamma}^{m} = \Sigma + W_{\beta\gamma} \quad (1)$$

**From contact angles to fluid-elastic parameters**

The contact angles depicted in Figure 3 are related to the distinct surface tensions along the three surface segments of the contact line (see Figures 3b-d). One of these surface tensions is the interfacial tension, $\Sigma_{\alpha\beta}$, between the condensate and the buffer. This interfacial tension balances the difference between the membrane tensions $\Sigma_{\alpha\gamma}^{m}$ and $\Sigma_{\beta\gamma}^{m}$ of the two membrane segments. This balance implies that the three surface tensions form the sides of a triangle (Figure 3c). Consequently, the mechanical tensions of the two membrane segments are described by Equation (1).

In the context of fluid-elastic parameters, $\Sigma$ represents the lateral stress within the membrane, conjugate to the total membrane area. The adhesion free energies per unit area, $W_{\alpha\gamma}$, $W_{\beta\gamma}$, and $W_{\alpha\beta}$, correspond to the interactions between the condensate (γ), the internal solution (β), and the external buffer (α), respectively. If the membrane prefers the condensate over the internal solution, the adhesion parameter $W_{\beta\gamma}$ is negative; conversely, if the internal solution is preferred, $W_{\beta\gamma}$ is positive. For simplification, the potential contribution of the spontaneous curvature of the membrane segments is neglected in Equation (1). The relative affinity between the condensate and the external buffer is given by:

$$W = \Sigma_{\beta\gamma}^{m} - \Sigma_{\alpha\gamma}^{m} = W_{\beta\gamma} - W_{\alpha\gamma} \text{ with} - \Sigma_{\alpha\beta} \leq W \leq + \Sigma_{\alpha\beta} \quad (2)$$

The inequality arises from the fundamental property of the tension triangle depicted in Figure 3, which states that the length of any side of a triangle must be less than or equal to the sum of the lengths of the other two sides. When the membrane prefers the condensate phase (γ) over the internal solution (β), the adhesion parameter $W_{\beta\gamma}$ is negative; conversely, when the internal solution is preferred, $W_{\beta\gamma}$ is positive. The limiting value $W=-\Sigma_{\alpha\beta}$ corresponds to complete wetting by the condensate phase, whereas the limiting case $W=+\Sigma_{\alpha\beta}$ represents dewetting from the condensate phase, which is equivalent to complete wetting by the external buffer. The tension triangle in Figure 3c also implies these relationships [2]:

$$\frac{\Sigma_{\alpha\gamma}^{m}}{\Sigma_{\alpha\beta}} = \frac{\Sigma + W_{\alpha\gamma}}{\Sigma_{\alpha\beta}} = \frac{sin\theta_{\beta}}{sin\theta_{\gamma}} \text{ and } \frac{\Sigma_{\beta\gamma}^{m}}{\Sigma_{\alpha\beta}} = \frac{\Sigma + W_{\beta\gamma}}{\Sigma_{\alpha\beta}} = \frac{sin\theta_{\alpha}}{sin\theta_{\gamma}} \quad (3)$$

According to the sine rule of the tension triangle in Figure 3c, the relationship between surface tension and contact angle is as follows. When subtracting the two equations in Equation (3), the relative affinity W in Equation (2) becomes equal to:

$$W = \Phi \equiv \frac{sin\theta_{\alpha} - sin\theta_{\beta}}{sin\theta_{\gamma}} \quad (4)$$

Therefore, the relative affinity $W=\Sigma_{\alpha\beta}$ is a mechanical quantity related to the adhesion free energy of the membrane segment, depending solely on the geometric factor Φ, which can be determined by analyzing the three contact angles obtained from optical images (Figure 3b). The inequality in Equation (2) implies that the geometric factor Φ satisfies: $1 \leq \Phi \leq 1$. The interpretation of the extreme values of Φ comes from the limiting cases of the affinity contrast W in Equation (2). $\Phi=-1$ corresponds to complete wetting of the membrane by the condensate phase, while $\Phi=+1$ corresponds to the condensate phase causing membrane dewetting. The dimensionless factor Φ is negative when the membrane prefers the condensate phase over the external buffer, and positive when the external buffer is preferred.

**Scanning Electron Microscope (SEM) characterization**

RAW264.7 cells were cultured on silicon wafers and incubated with DNA droplets for 1 hour at 37°C. Subsequently, the cells were fixed overnight at 4°C using a solution containing 2% paraformaldehyde, 2.5% glutaraldehyde, and 0.1 M sodium cacodylate. Following fixation, the samples were dehydrated through a graded ethanol series. Critical point drying was performed using $CO_2$, and the samples were then gold-coated. Imaging was conducted using a field emission scanning electron microscope (10-15 keV, JSM-7500F, JEOL).

**Plant culture**

Seeds of wild-type *Nicotiana benthamiana* were surface-sterilized and placed onto solidified culture medium in a growth room (20-22 °C day/16-18 °C night, 12 h/12 h). After seedlings had developed leaves, the leaves were excised and transferred to a callus induction medium for the cultivation of callus. The induction medium contains Murashige and Skoog (MS) basal salts, plant growth regulators (1mg/mL 2,4-D), sucrose, and agar, with the medium adjusted to a pH of 5.7.

**Protoplast isolation**

Four-week-old *Nicotiana benthamiana* leaves were harvested and thin leaf strips were prepared using a fresh, sharp razor blade, ensuring that the tissue was not crushed. The leaf strips were transferred to a filtered enzyme solution (approximately 2 leaves in 5 mL of enzyme solution). The enzyme solution contained 20 mM MES, 1.5% (w/v) cellulase R10 (Duchefa Biochemie), 0.4% (w/v) macerozyme R10 (Duchefa Biochemie), 0.4 M mannitol, 20 mM KCl, and was adjusted to pH 5.7. The digestion process was carried out at room temperature in the dark with gentle shaking for at least 3 hours. The enzyme/protoplast solution was then diluted with an equal volume of MMG solution (containing 0.4 M mannitol, 15 mM $MgCl_2$, and 4 mM MES, pH 5.7). Prior to use, a clean nylon mesh (75 μm) was soaked in 95% ethanol, washed with sterile water, and wetted with MMG solution. The enzyme solution containing protoplasts was then passed through the nylon mesh to remove undigested leaf material. The filtrate was centrifuged for 2 minutes at 50 mL in a round-bottom tube to pellet the protoplasts. After removing the supernatant, the protoplast pellet was gently resuspended in MMG solution by gentle rotation. The protoplasts were allowed to settle by gravity at the bottom of the tube for 15 minutes. The supernatant was removed, and the protoplasts were resuspended in MMG solution to a final concentration of approximately $10^5$ cells $mL^{-1}$. For the callus tissue protoplast extraction, the procedure was the same as for the leaves, but the enzymatic digestion time was extended to 6 hours.

**Liquid chromatograph-mass spectrometer (LC-MS) characterization**

Lipid extraction and sample preparation were performed following the method described in the referenced literature [3]. Liquid chromatography-tandem mass spectrometry (LC-MS/MS) analysis was carried out using the Shimadzu LC-20AD system (Kyoto, Japan), coupled with the Sciex X500R QTOF mass spectrometer (Toronto, Canada). Sample processing was conducted by Major Biomedical Technology (Shanghai, China), and the resulting data were analyzed using SciPy (Python) Version 1.0.0.

**Cellular Uptake of DNA droplets**

RAW264.7 cells were seeded in a CLSM dish and incubated overnight. Subsequently, Cy3-

labeled DNA droplets were added to the culture medium at a final concentration of 100 nM and incubated for 12 hours. After three washes with 1× PBS, the cells were stained with CellMask™ plasma membrane stain for 10 minutes. Imaging was performed using CLSM to observe the cellular uptake of the DNA droplets. Quantitative analysis of the mean fluorescence intensity of CLSM images was performed using Image J.

**Evaluation of uptake mechanisms via endocytosis inhibition**

RAW264.7 cells were seeded in CLSM dishes and incubated overnight. After washing with 1× PBS, the cells were incubated for 1 hour at 37°C in a 5% $CO_2$ atmosphere with culture medium containing endocytosis inhibitors: cytochalasin D (20 μg/mL), chlorpromazine (10 μg/mL), methyl-β-cyclodextrin (10 mmol/L), or dynasore (100 μmol/L). Following treatment, the cells were washed with 1× PBS, and pre-constructed Cy3-labeled DNA droplets (with a final concentration of 100 nM) were added to the dishes and incubated for 12 hours. After three washes with 1× PBS, the cells were stained with CellMask™ plasma membrane stain and observed using CLSM.

**Loading of DNA droplets with functional molecules**

Typically, the oligodeoxynucleotide cytidylyl phosphate guanosine (CpG) is directly incorporated into DNA droplets through sequence design. Streptavidin (SA) is introduced by specific binding to biotinylated DNA strands (S1-biotin in Table S1). Luciferase mRNA is incorporated by mixing mRNA (1 μg/μL) with DNA monomers at a 1:10 ratio (mRNA polyA to DNA monomer polyT) in 1× PBS, followed by incubation at 37°C for 1 hour, facilitating effective mRNA enrichment during droplet formation.

**Gel electrophoresis**

The mRNA and DNA droplet complexes were loaded onto a 2% agarose gel, with 1× loading buffer added to each sample. Electrophoresis was performed in 1× TAE buffer (40 mM Tris-acetate, 1 mM EDTA, pH 8.0) at 80 V under ice-cold conditions. The electrophoresis was conducted using a Bio-Rad electrophoresis system.

**Induction of cytokine secretion by CpG-functionalized DNA droplets**

RAW264.7 cells were seeded in 24-well plates at a density of $1\times10^5$ cells per well and incubated for 12 hours. Prior to treatment, cells were washed twice with PBS. DNA droplets were diluted to a final concentration of 100 nM in culture medium and added to the cells. The cells were incubated at 37°C for 8 to 72 hours, after which the culture supernatants were collected for TNF-α analysis. TNF-α levels in the supernatants were measured using an enzyme-linked immunosorbent assay (ELISA) kit, following the manufacturer's instructions.

**Delivery of Luciferase mRNA**

RAW264.7 or plant cells (protoplasts derived from callus tissue) were seeded in 96-well plates and incubated overnight to allow attachment to the glass bottom. Luciferase mRNA was delivered either as free mRNA or encapsulated within DNA droplets. In both cases, the amount of mRNA was 1 μg per well. After a 48-hour incubation, cells were collected, and luciferase activity was measured using a luciferase reporter assay system (Promega, Madison, USA).

**Table S1. Sequences of oligonucleotides used in this study.**

| Name | Sequences (from 5' to 3') |
|---|---|
| **S1** | CGCGAGCAAACACGGCTACGAATCGAATTCCCGTTGGC |
| **S2** | CGCGCTGACATCCGCTTCGGAACGTAGCCGTGTTTGCT |
| **S3** | CGCGTAGCACAGAGGCTCGCAACCGAAGCGGATGTCAG |
| **S4** | CGCGGCCAACGGGAATTCGAAAGCGAGCCTCTGTGCTA |
| **Chol-S2** | Chol-AACTGACATCCGCTTCGGAACGTAGCCGTGTTTGCT |
| **S3-Cy3** | CGCGTAGCACAGAGGCTCGCAACCGAAGCGGATGTCAG-Cy3 |
| **S4-Chol** | CGCGGCCAACGGGAATTCGAAAGCGAGCCTCTGTGCTA-Chol |
| **Five-S1** | CGCGGTGCTATGAGGTTGCGTTCCTGGTACTGTACATG |
| **Five-S2** | CGCGCATGTACAGTACCAGGTTGCATCGATAGTCGACG |
| **Five-S3** | CGCGCGTCGACTATCGATGCTTGCCTTGACTGCCAATC |
| **Five-S4** | CGCGGATTGGCAGTCAAGGCTTCGAAGTTGGAGCAGAC |
| **Five-S5** | CGCGGTCTGCTCCAACTTCGTTCGCAACCTCATAGCAC |
| **Five-S1-Cy3** | CGCGGTGCTATGAGGTTGCGTTCCTGGTACTGTACATG-Cy3 |
| **Five-Chol-S2** | Chol-CATGTACAGTACCAGGTTGCATCGATAGTCGACG |
| **Five-S5-Chol** | CGCGGTCTGCTCCAACTTCGTTCGCAACCTCATAGCAC-Chol |
| **S1-CpG** | CGCGAGCAAACACGGCTACGAATCGAATTCCCGTTGGCTCCATGACGTTCCTGACGTT |
| **S1-CpG-FAM** | CGCGAGCAAACACGGCTACGAATCGAATTCCCGTTGGCTCCATGACGTTCCTGACGTT-FAM |
| **PolyT30-S1** | TTTTTTTTTTTTTTTTTTTTTTTTTTTTTTAGCAAACACGGCTACGAATCGAATTCCCGTTGGC |
| **S1-biotin** | CGCGAGCAAACACGGCTACGAATCGAATTCCCGTTGGC-biotin |
| **S1-free** | AGCAAACACGGCTACGAATCGAATTCCCGTTGGC |
| **S2-free** | CTGACATCCGCTTCGGAACGTAGCCGTGTTTGCT |
| **S3-free** | TAGCACAGAGGCTCGCAACCGAAGCGGATGTCAG |
| **S4-free** | GCCAACGGGAATTCGAAAGCGAGCCTCTGTGCTA |

**Table S2. Fluid properties of DNA droplets used in this study.**

| Droplet | FRAP recovery | $t_{1/2}$ (min) | η/γ (min / μm) |
|---|---|---|---|
| **4SE** | 95% | 3.52 | 6.94 |
| **Chol'-4SE** | 75% | 4.06 | 8.02 |
| **Chol-4SE** | 46% | 13.47 | 26.68 |
| **Chol-3SE** | 65% | 6.26 | 13.92 |
| **2Chol'-3SE** | 52% | 7.13 | 17.43 |

### Supporting Figures

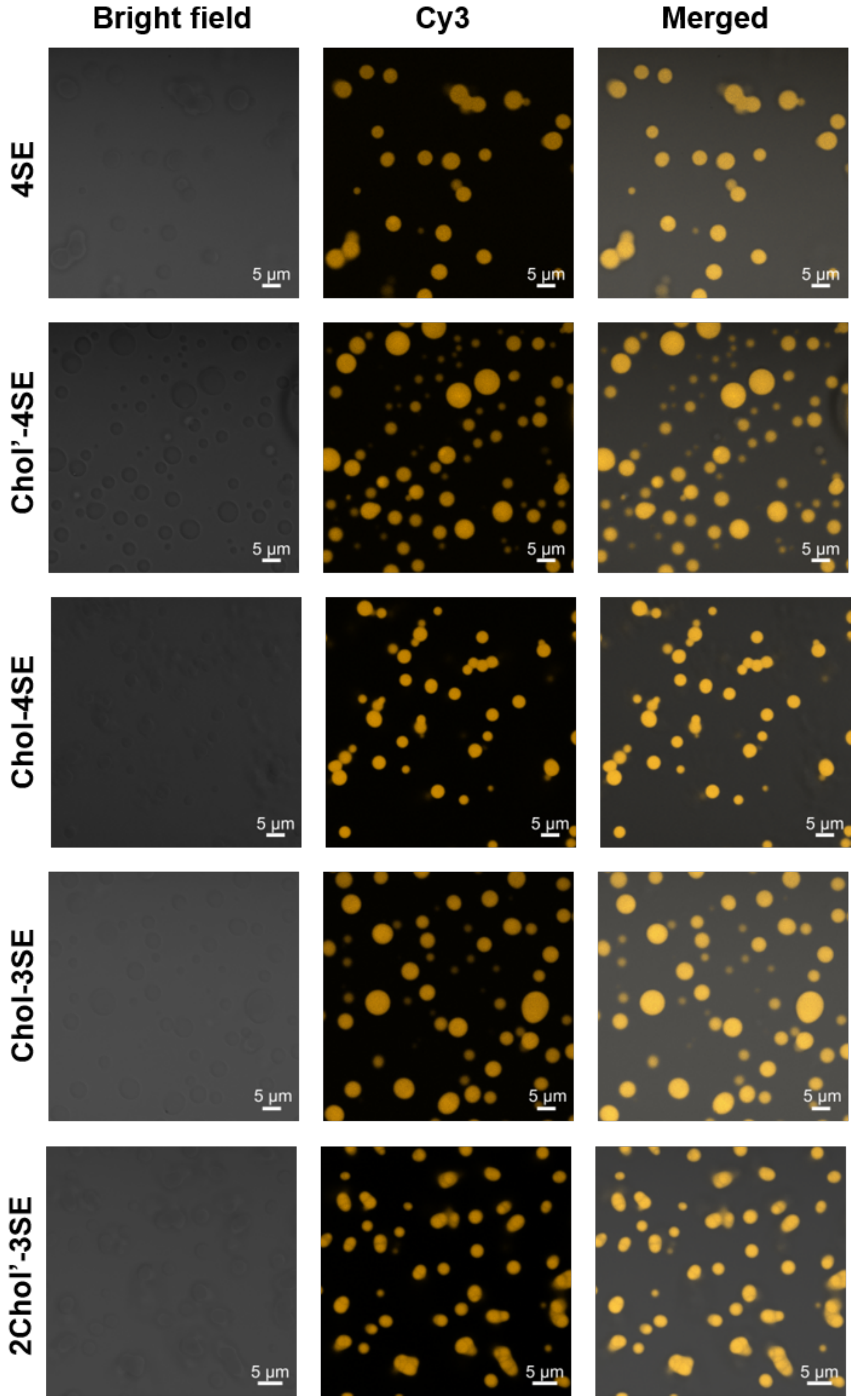


**Figure S1.** The representative CLSM images of DNA droplets formed by different DNA monomers (4SE, Chol'-4SE, Chol-4SE, Chol-3SEand 2Chol'-3SE). Scale bars: 5 μm.

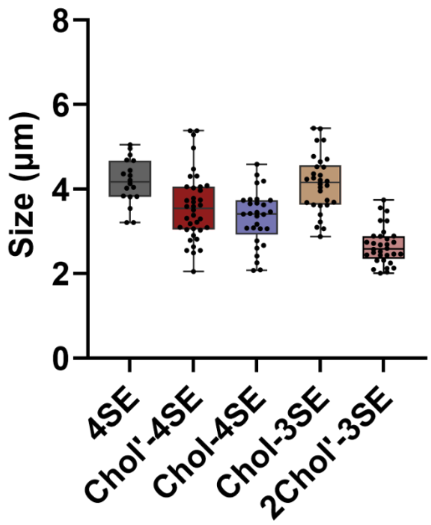


**Figure S2.** The statistical analysis of size distribution of droplets formed by various DNA monomers.

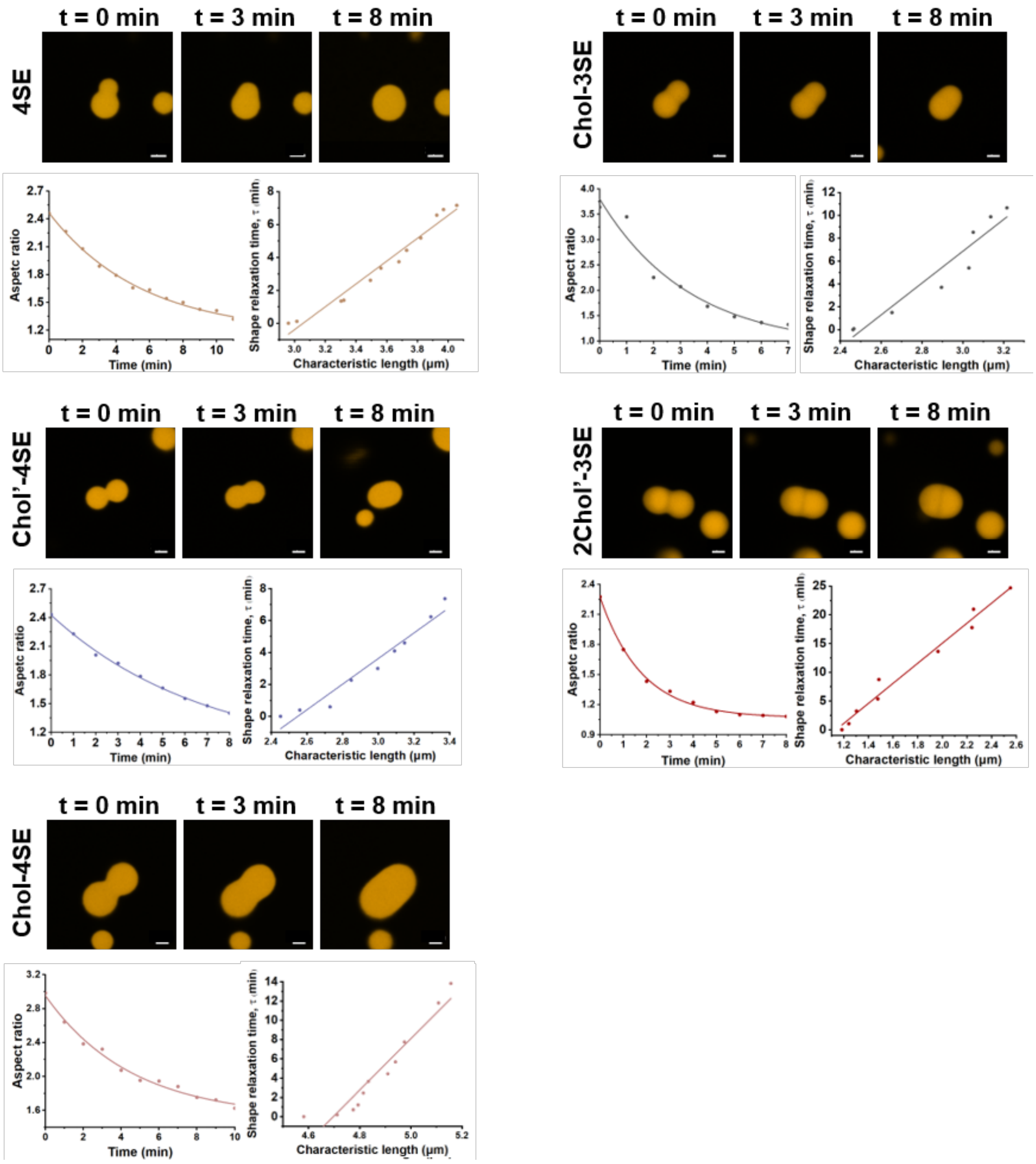


**Figure S3.** Top panel: CLSM images of the fusion process of different DNA droplets in 1× PBS. Bottom panel: time-dependent changes in the aspect ratio (*AR*) of droplets during the fusion process. Data were fit with an exponential function to extract the shape relaxation timescale ($\tau$). Relationship between shape relaxation time ($\tau$) and droplet length scale ($l$). The slope of the linear fit corresponds to the inverse capillary velocity ($\eta/\gamma$). All experiments were conducted at room temperature. Scale bars: 2 μm.

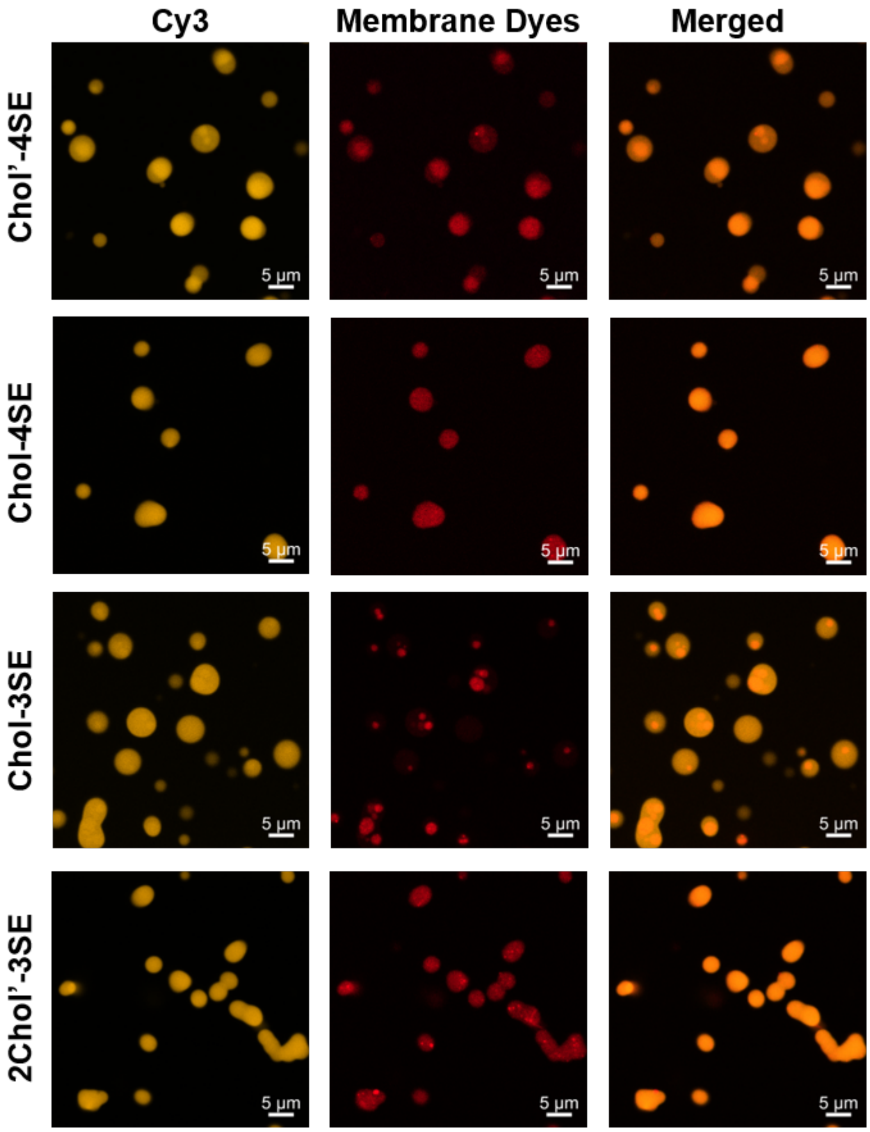


**Figure S4.** Representative confocal images showing the distribution of cholesterol molecules within different DNA droplets. Cy3-labeled DNA droplets are shown in yellow, and cholesterol is visualized using a lipid-specific fluorescent dye (red). Scale bars: 5 μm.

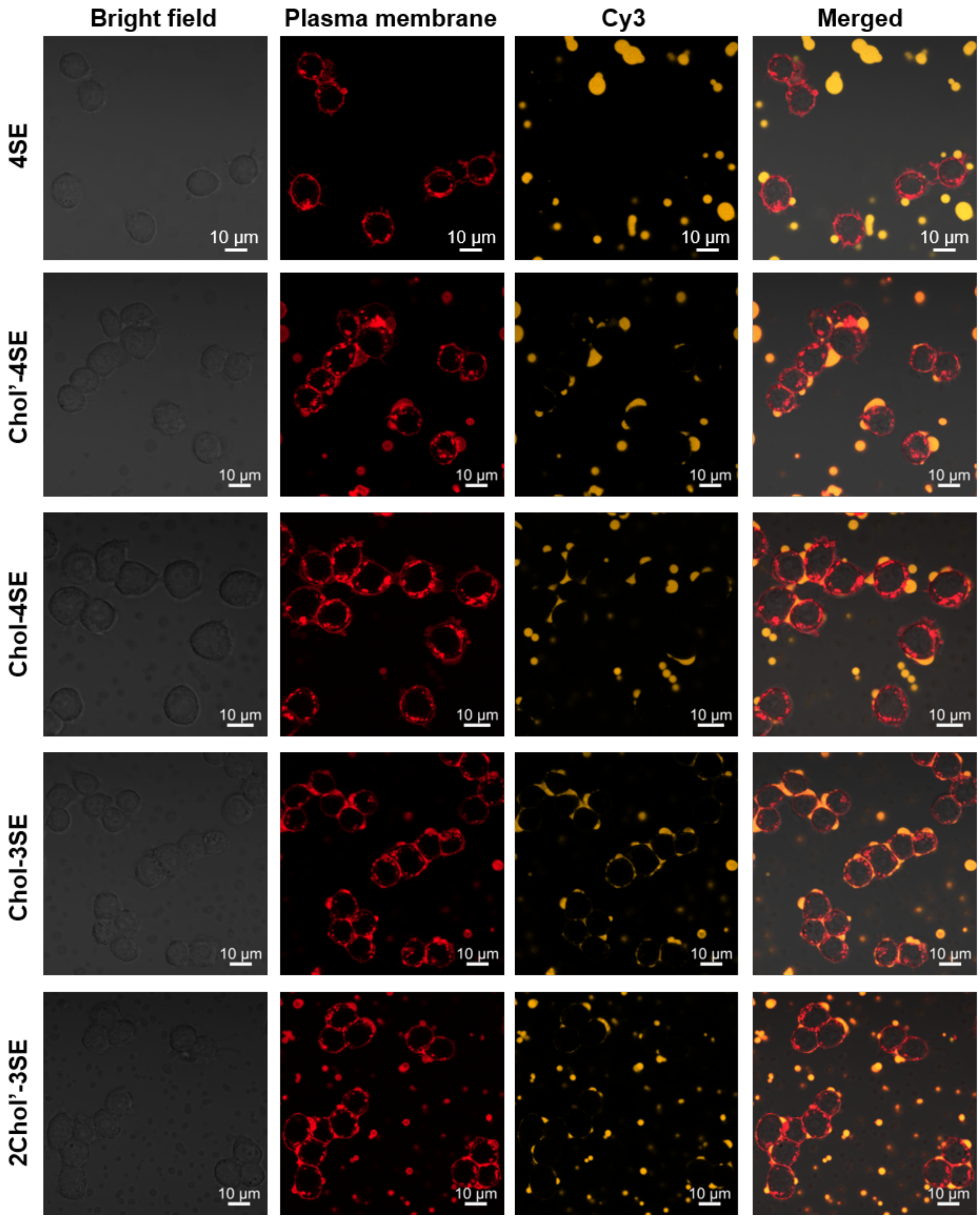


**Figure S5.** Representative CLSM images showing the interaction between plasma membrane labeled with lipid dye CellMask™ (red) and Cy3-labeled TDC droplets (orange) (1×PBS, 1 μM TDC droplets, 37 °C, 2 hours incubation). Scale bars: 10 μm.

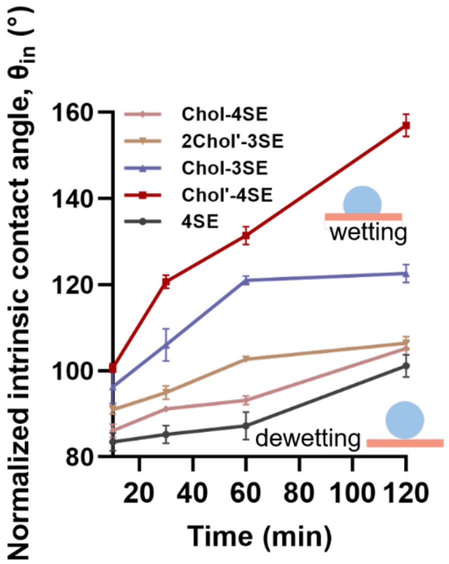


**Figure S6.** Analyzing the wetting behaviors of TDC droplets using the contact angel. The contact angles were measured from confocal images of DNA droplets, with data points representing the mean values from three independent experiments.

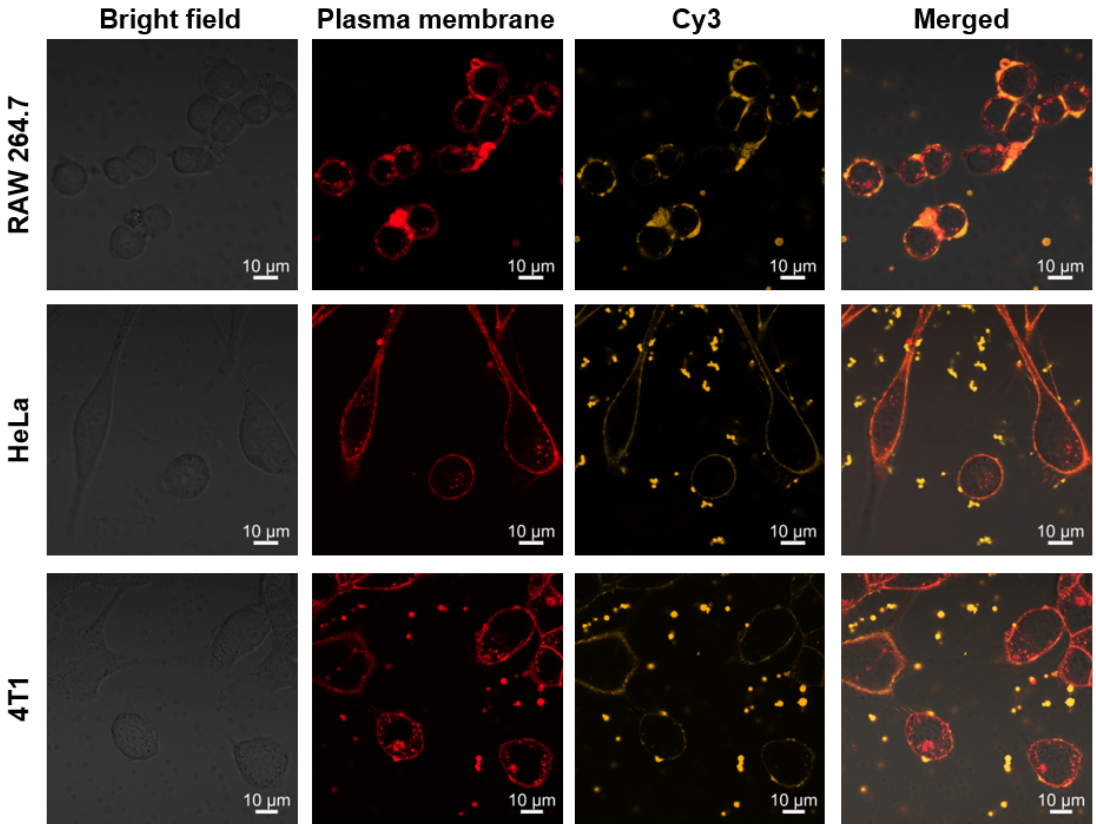


**Figure S7.** Representative CLSM images showing the interaction between Chol-3SE droplet and various cells with different cholesterol contents. The cell membranes were labeled with lipid dye CellMask™ (red) and TDC droplets were Cy3-labeled (orange) (1×PBS, 1 μM droplets, 37 °C, 2 hours incubation). Scale bars: 10 μm.

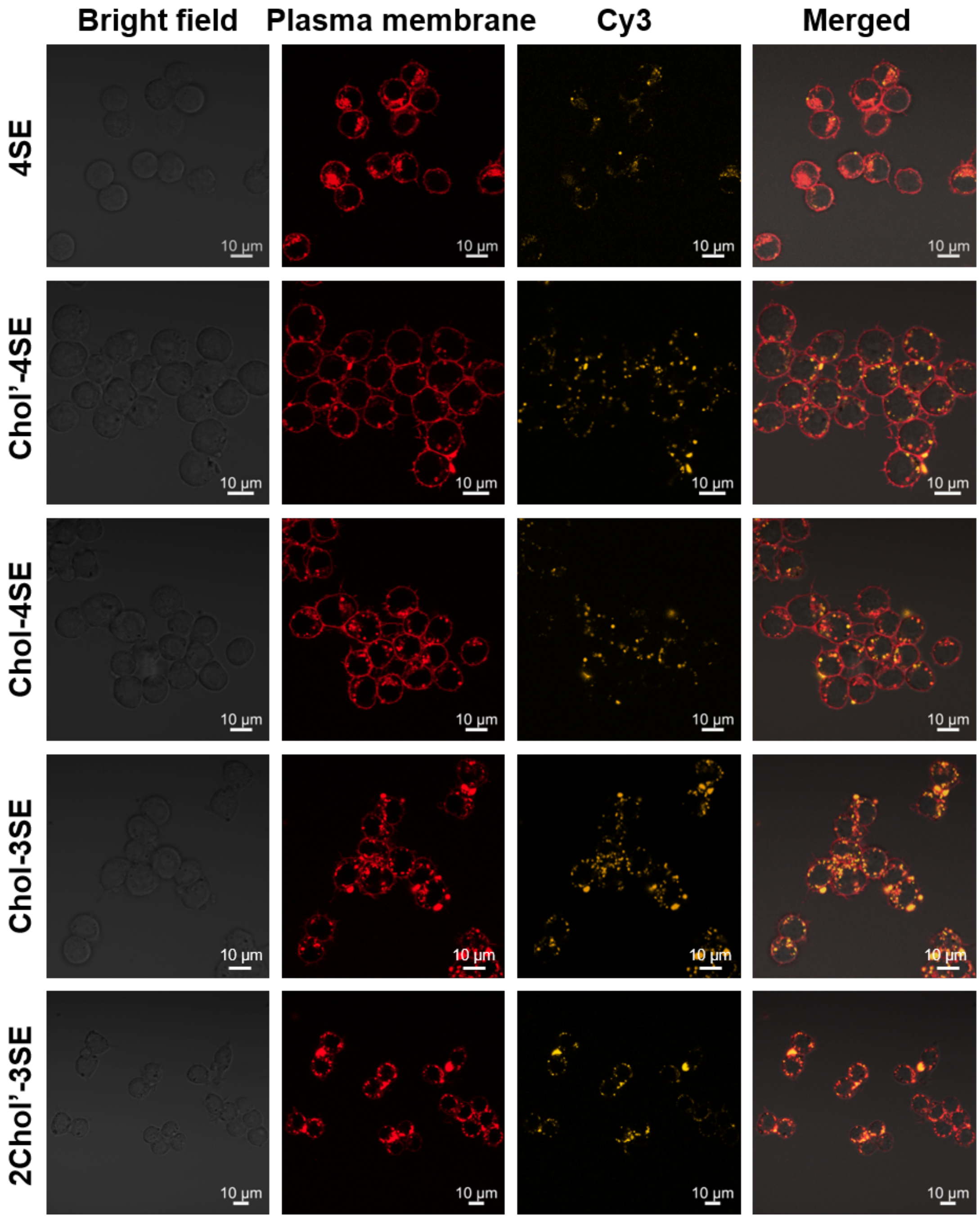


**Figure S8.** Representative CLSM images showing the internalization of various TDC droplets by RAW 264.7 cells after 12h incubation. Scale bars: 10 μm.

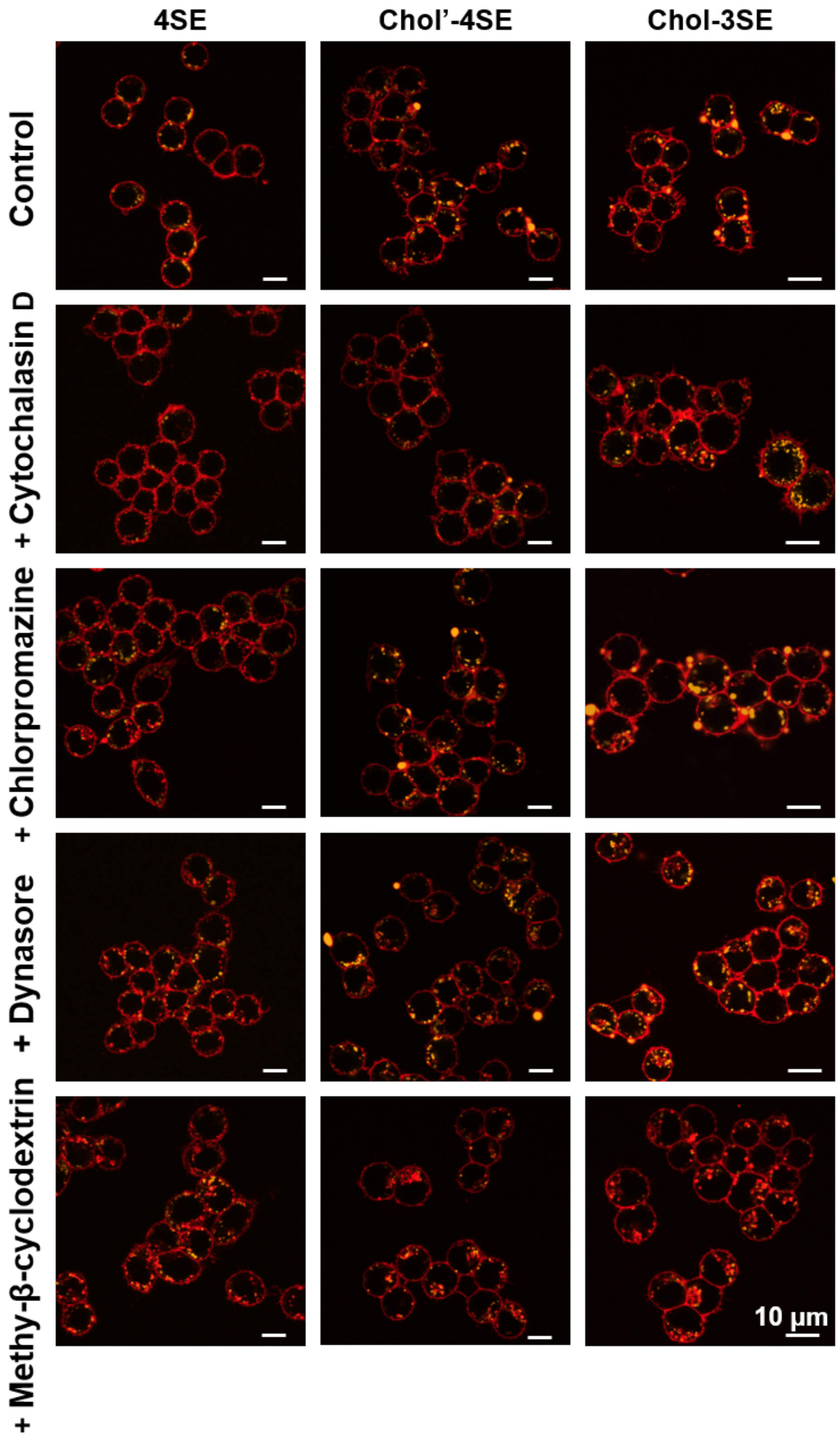


**Figure S9.** Representative CLSM images of untreated RAW264.7 cells and RAW264.7 cells treated with different uptake inhibitors. The inhibitors used include Cytochalasin D, Chlorpromazine, Dynasore, and Methyl-β-cyclodextrin. Scale bars: 10 μm.

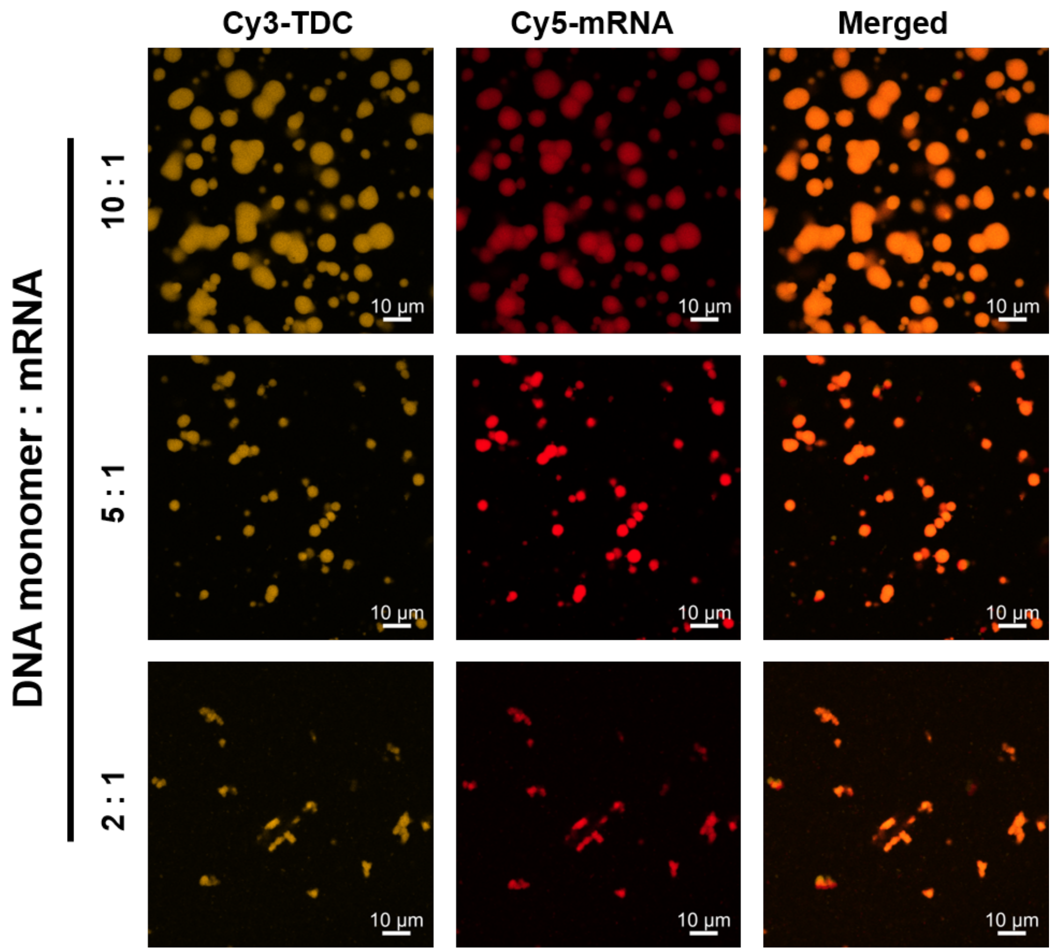


**Figure S10.** Representative CLSM images of Chol'-4SE droplets formed by different molar ratio of polyT-DNA monomer and polyA-mRNA. Scale bars: 10 μm.

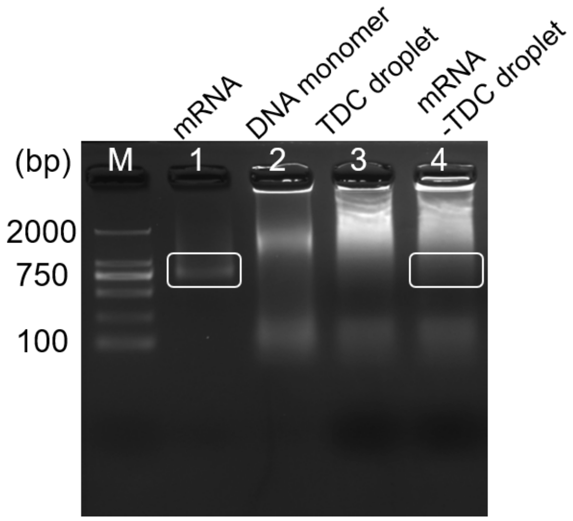


**Figure S11.** 2% agarose gel characterization of Luciferase mRNA loading within the droplet under the condition of mRNA: DNA monomer ratio of 1:2. Lane 1: Luciferase mRNA (~2000 nt). Lane 2 and Lane 3: the Chol-DNA monomer and Chol-DNA droplet. Lane 4: mRNA loaded DNA droplet (Chol'-4SE). M: marker.

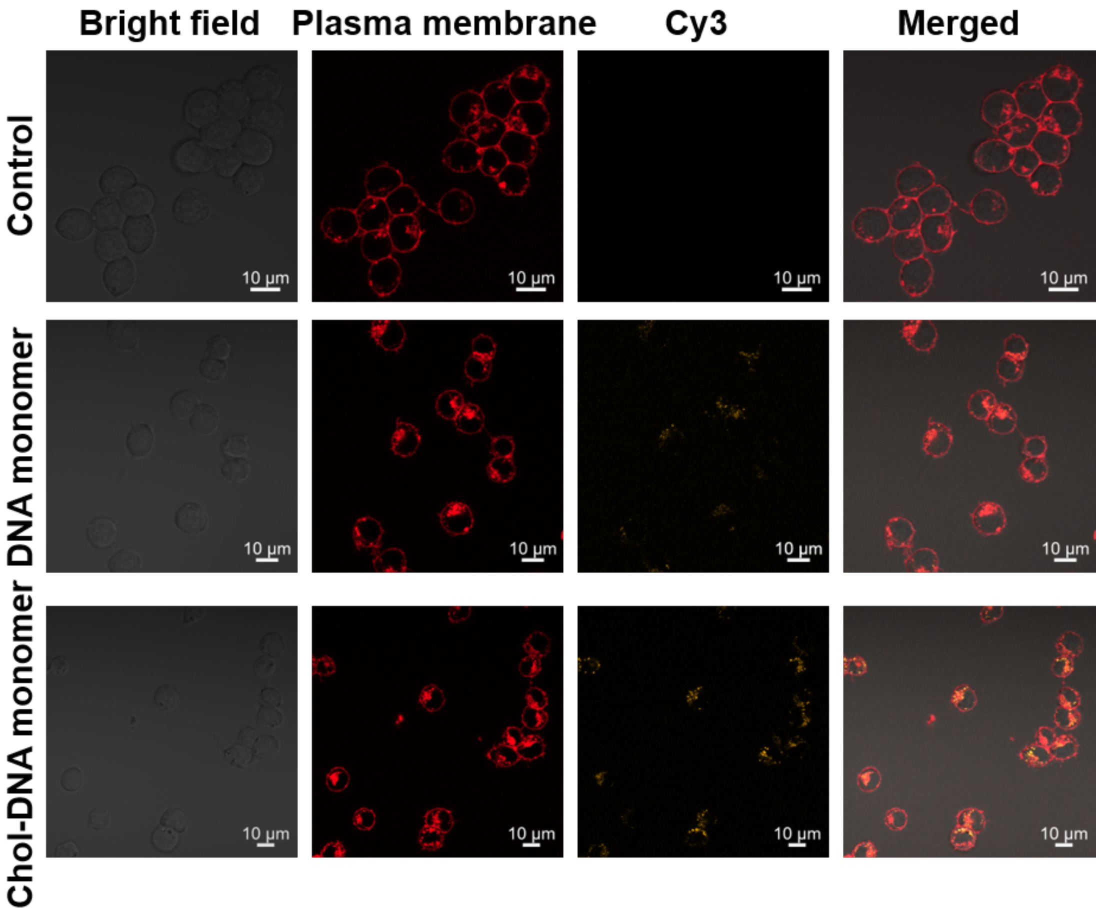


**Figure S12.** CLSM images showing the internalization of DNA monomer or Chol-DNA monomer into RAW264.7 cells after 12h incubation. Scale bars: 10 μm.

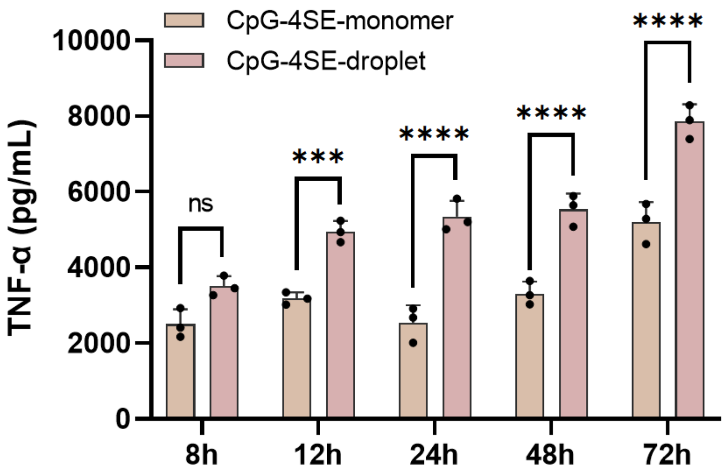


**Figure S13.** RAW 264.7 cells were stimulated with free-CpG, CpG-DNA monomer, or CpG-DNA droplets (4SE) at a final CpG concentration of 100 nM, and secretion of the pro-inflammatory cytokine TNF-α was assessed (n = 3 biological replicates). Statistical significance is indicated as follows: ***, $P < 0.001$; ****, $P < 0.0001$, one-way ANOVA.

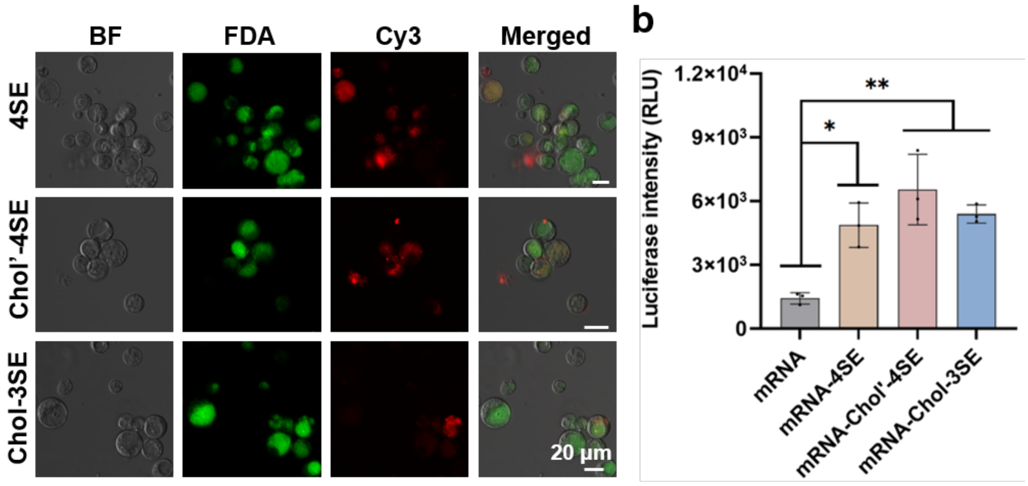


**Figure S14.** (a) Representative CLSM images showing uptake of various TDC droplets by protoplast (24 hours incubation). Protoplast was stained with FDA (green), and droplets were Cy3-labeled (red). Scale bars: 20 µm. (b) Luciferase activity evaluation after 48 hours incubation with free mRNA or mRNA loaded TDC droplets (n = 3 biological replicates).

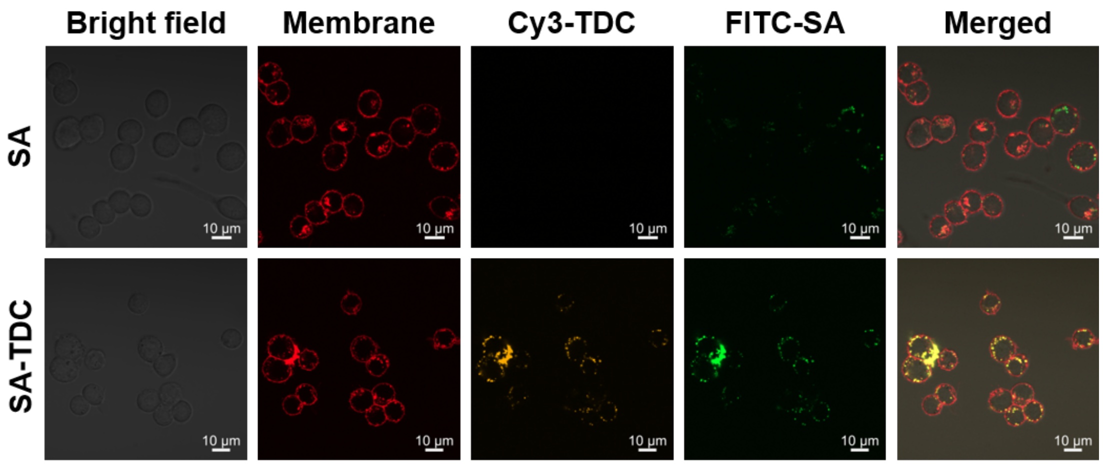


**Figure S15.** CLSM images show the internalization of free-SA or SA-loaded Chol'-4SE droplets by RAW 264.7 cells after 12h incubation. Scale bars: 10 µm.

**Movie S1-S5.** Time-lapse videos (0-2 hours) of various DNA droplets: 4SE (Movie S1), Chol'-4SE (Movie S2), Chol-4SE (Movie S3), Chol-3SE (Movie S4), and 2Chol'-3SE (Movie S5), incubated with RAW264.7 cells were captured using Confocal Laser Scanning Microscopy (CLSM). Scale bars: 2 µm or 5 µm.
